# Synaptic Organization-Function Relationships of Amygdala Interneurons Supporting Associative Learning

**DOI:** 10.1101/2024.06.18.599631

**Authors:** Rita Báldi, Sunil Muthuswamy, Niharika Loomba, Sachin Patel

## Abstract

Coordinated activity of basolateral amygdala (BLA) GABAergic interneurons (INs) and glutamatergic principal cells (PCs) is critical for associative learning, however the microcircuit organization-function relationships of distinct IN classes remain uncertain. Here, we show somatostatin (SOM) INs provide inhibition onto, and are excited by, local PCs, whereas vasoactive intestinal peptide (VIP) INs are driven by extrinsic afferents. Parvalbumin (PV) INs inhibit PCs and are activated by local and extrinsic inputs. Thus, SOM and VIP INs exhibit complementary roles in feedback and feedforward inhibition, respectively, while PV INs contribute to both microcircuit motifs. Functionally, each IN subtype reveals unique activity patterns across fear- and extinction learning with SOM and VIP INs showing most divergent characteristics, and PV INs display an intermediate phenotype parallelling synaptic data. Finally, SOM and PV INs dynamically track behavioral state transitions across learning. These data provide insight into the synaptic microcircuit organization-function relationships of distinct BLA IN classes.

## INTRODUCTION

Associative learning facilitates survival by optimizing behavioral responses to changing environmental contexts and cues ^1,2^. These learned behavioral responses promote both the securing of resources needed for survival and reproduction and the avoidance of real or potential threats. Importantly, the expression of learned defensive responses to environmental cues must be balanced with the physiological extinction of these responses when the associations between environmental cues and danger no longer exist ^3^. Failure to appropriately extinguish conditioned defensive responses may compromise behaviors aimed at resource and reward acquisition. Extinction impairment also serves as a hallmark of several neuropsychiatric disorders, including posttraumatic stress disorder (PTSD) ^4,5^. Thus, elucidating the mechanisms governing aversive associative learning processes could provide valuable insight into the pathophysiology of stress and trauma-related disorders.

The amygdala is well-established as a key site for associative learning and synaptic plasticity within the circuits contributing to aversive associative learning and extinction ^6–8^. Indeed, distinct basolateral amygdala (BLA) pyramidal cell (PC) populations defined by projection target or molecular phenotype have been implicated in both acquisition and extinction of conditioned fear behavior using rodent models ^9–12^. More recently, the role of specific interneuron (IN) subtypes has been investigated in aversive associative learning. Similar to other cortical-like structures, the BLA contains largely non-overlapping populations of GABAergic INs expressing somatostatin, parvalbumin, and vasoactive intestinal peptide (SOM, PV, VIP), that have been implicated in the regulation of associative learning^13–16^. For instance, using an auditory fear conditioning paradigm, investigators observe decreased SOM IN activity following conditioned stimulus (CS) presentation, contributing to enhanced cue-shock associations^17^. Conversely, increased SOM IN activity is associated with reduced fear behavior and theta frequency oscillations in the presence of learned safety signals ^18^. VIP INs are activated by aversive unconditioned stimulus (US; shock) and strongly modulated by expectation, and mediate adaptive disinhibitory gating supporting associative learning ^19^. Similarly, PV INs are US responsive in an expectancy-dependent manner across CS-US pairings ^20^, although another study found suppression following US deliveries ^17^. Furthermore, enhanced CS-induced PV IN activity could promote fear learning and retrieval via the inhibition of SOM INs ^17^, and disinhibition of PCs. From a synaptic connectivity perspective, VIP INs are reported to primarily inhibit the activity of PV and SOM INs in the BLA, while SOM and PV INs exert inhibition over PCs ^19^. These data highlight critical roles for BLA INs in the regulation of aversive associative learning and suggest that a comprehensive analysis of BLA IN microcircuit connectivity and in vivo activity patterns could provide further clarification into the mechanisms underlying fear memory formation and subsequent extinction.

Despite the aforementioned data, a variety of critical questions related to the roles and organization of BLA INs remain unanswered. These include, how distinct INs are regulated by extrinsic long-range inputs and intrinsic BLA PCs, how INs gate extrinsic afferent-induced activation of BLA PCs, how different IN subtypes represent US and CS information during fear acquisition and extinction, and how IN activity relates to the expression of freezing behavior. To address these questions, we show that distinct IN subtypes contribute to different synaptic microcircuit motifs, with SOM and VIP INs participating in feedback and feedforward inhibition, respectively, and PV INs contributing to both. In vivo, we identify divergent activity patterns for SOM and VIP INs across fear conditioning and extinction, with PV INs displaying intermediate phenotypes with properties of both SOM and VIP INs. Moreover, we find that SOM and PV INs dynamically track behavioral state transitions between motion and freezing across fear acquisition and extinction. Altogether, these data demonstrate that synaptic organization-function relationships exist, wherein INs generating feedforward or feedback inhibition distinctively and dynamically represent threat-predictive environmental stimuli and behavioral state transitions as a function associative learning.

## RESULTS

### Differential Long-Range Targeting of BLA IN subtypes

To begin to understand potential differential synaptic regulation of BLA IN activity we first examined spontaneous excitatory postsynaptic currents (sEPSCs) onto BLA PCs and SOM, PV, and VIP INs using whole-cell voltage-clamp recordings (**Fig. S1 a-b**). SOM and PV INs exhibited higher sEPSC frequency than PCs, suggesting SOM and PV INs may receive distinct excitatory inputs relative to PC and VIP INs. sEPSC amplitude was also larger in SOM and PV INs relative to PCs indicating diversity in postsynaptic efficacy. Furthermore, varying differences in inhibitory transmission were observed between cell types (**Fig. S1 c-d**). These data suggest that specific IN populations are innervated by differing excitatory afferents; thus, we sought to uncover the potentially unique long-range connectivity between limbic forebrain structures and BLA IN subtypes.

Using SOM, PV, and VIP reporter mice, we first expressed an excitatory opsin in areas implicated in associative learning ^21–23^ (prelimbic cortex (PL), dorsal midline thalamus (DMT), or lateral entorhinal cortex (LEC)) and conducted whole-cell current-clamp recordings from pairs of identified INs and neighboring PCs (**Fig. 1 a-b**). Excitatory postsynaptic potentials (EPSPs) in PCs were monophasic, while SOM and PV INs often showed multiphasic responses with two identified peaks (**Fig. 1 c**). Specifically, upon PL stimulation, all SOM and PV, while only 13 % of VIP INs, showed long-latency responses (**Fig. 1 d-e**). Importantly, long-latency responses were much larger in SOM and PV INs than monophasic EPSPs (**Fig. 1 d and f**). Application of TTX+4AP confirmed short-latency first peaks in SOM, PV, and VIP INs to be monosynaptic as no significant change in amplitude was noted relative to baseline EPSP amplitudes, while the large amplitude long-latency responses were abolished verifying their disynaptic nature (see **Fig. 1 c and f**). Following DMT terminal stimulation 100% of SOM, 57% of PV, and 40% of VIP INs showed disynaptic long latency responses, with disynaptic responses being larger in SOM INs (**Fig. 1 g-h**). Short-latency EPSPs in SOM and PV INs remained comparable after TTX+4AP application with no second peaks present, validating monosynaptic phenotype (**Fig. 1 i**). Upon LEC stimulation, 91% of SOM, but only 20% of PV and 22% of VIP INs, exhibited disynaptic responses, and again, these disynaptic peaks were significantly larger than the monosynaptic EPSPs in SOM INs, and were eliminated after the application of TTX+4AP (**Fig. 1 j-l**). These data demonstrate that PL, DMT, and LEC afferents can excite all IN subtypes in the BLA and can trigger disynaptic responses to varying degrees depending on the input stimulated and the IN type. Importantly, SOM INs appear to receive more robust disynaptic excitation, both in terms of number of cells demonstrating disynaptic responses and the overall amplitude of these disynaptic EPSPs, relative to PV and VIP INs. In contrast, VIP INs generally show the smallest degree of disynaptic responses and receive overall the strongest monosynaptic inputs from the PL and DMT. PV INs show an intermediate or mixed phenotype, resembling SOM INs upon PL stimulation, but VIP INs upon DMT stimulation, for instance.

**FIGURE 1.**
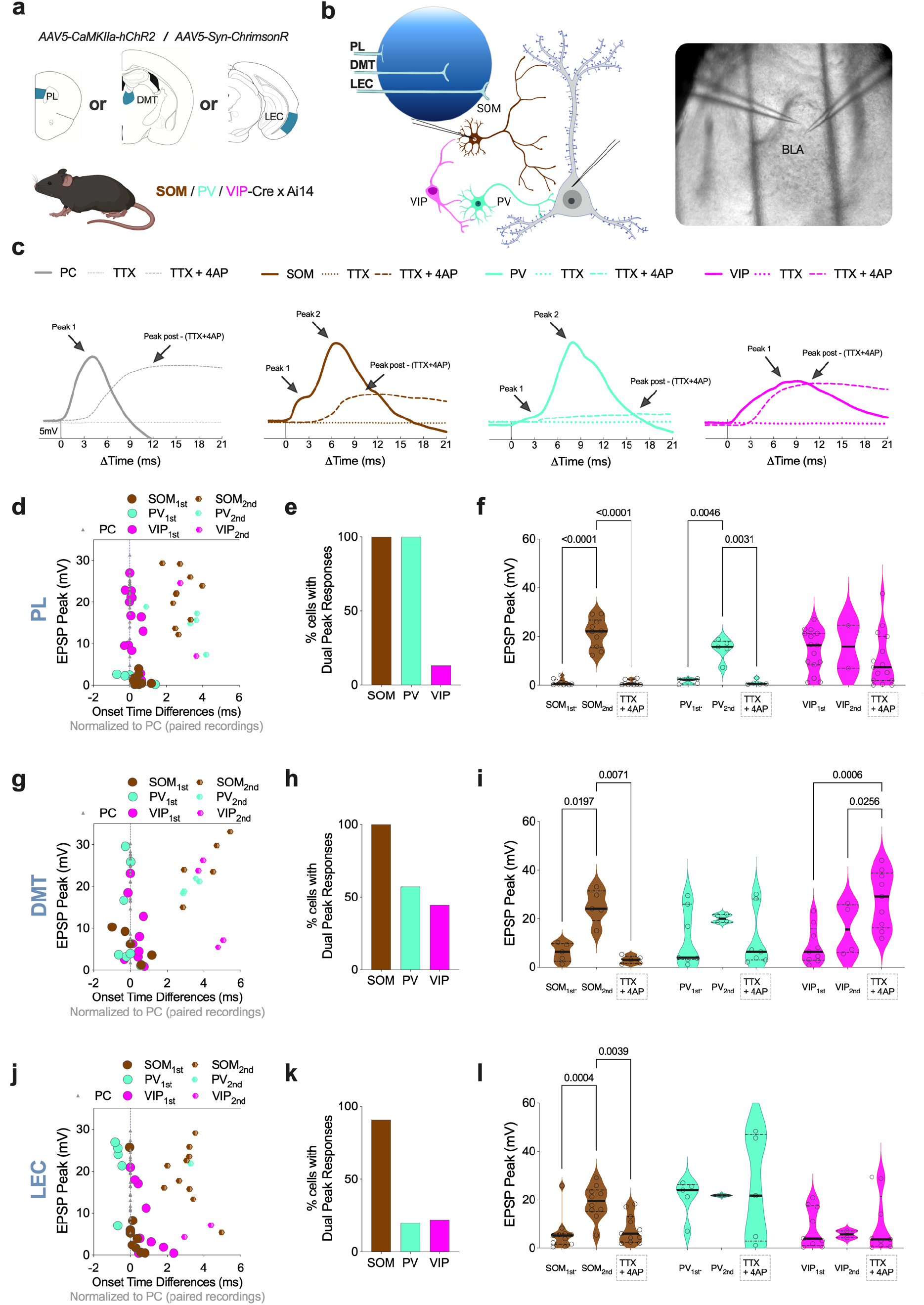
Distinct long-range connectivity patterns across BLA IN types. a-b, Diagrams of experimental design. An excitatory opsin encoding AAV was injected into the PL, DMT, or LEC of SOM-, PV- or VIPxAi14 mice to perform whole-cell current clamp recordings of adjacent BLA PC and IN pairs. **c,** Representative traces of cell-types upon single stimulation at baseline, after TTX, and after TTX+4AP drug applications. **d-f,** Data derived from PL input stimulation. **d,** X-Y plot of EPSP peaks and EPSP onset time differences relative to the paired PC of individual SOM, PV, and VIP INs. **e,** Proportion of cells exhibiting long-latency disynaptic responses per IN types. **f,** Quantification of observed monosynaptic (_1st_), disynaptic (_2nd_) and confirmed monosynaptic (TTX+4AP) EPSP peaks for each IN type. **g-i,** Data parallel to **d-f** upon DMT terminal stimulation. **j-l,** Data parallel to **d-f** following LEC input stimulation. Sample sizes of PC-IN pairs (afferent: IN type (n=cell pairs, number of mice)); PL: SOM (10,4), PV (5,3), VIP (15,6); DMT: SOM (5,2), PV (7,3), VIP (9,4); LEC: SOM (11,5), PV (5,2), VIP (9,4). **f**, **i**, **l,** Statistical analysis and P values via repeated measures two-way ANOVA followed by Tukey post hoc analysis (see **Table S1**).

### Differential Excitation of BLA INs from Extrinsic and Intrinsic Inputs

Our data suggest SOM and PV INs receive substantial disynaptic excitation upon long-range afferent stimulation, relative to VIP INs. Since the only source of local excitation to BLA INs are BLA PCs, these findings imply that SOM and PV INs receive more robust internal excitation from local PCs than from extrinsic sites. To test this hypothesis, we examined the ability of extrinsic afferents, from the PL, DMT and LEC, as well as intrinsic inputs from local BLA PCs, to elicit action potential firing of BLA INs and neighboring PCs. To isolate extrinsic afferents, we injected an excitatory opsin encoding AAV into the PL, DMT, or LEC of SOM, PV, or VIP IN reporter mice and performed paired recordings of INs and adjacent PCs (**Fig. 2 a**). To isolate local intrinsic inputs to BLA INs, we injected a Flpo-recombinase encoding retrograde-AAV into the PL or the lateral nuclei of central amygdala (CeL) of SOM-, PV-, or VIP-Ai14 mice, in addition to the combination of Cre-off/Flp-on ChRmine and DIO-eYFP encoding viruses into the BLA to perform electrophysiological recordings from non-labeled PCs and eYFP-labeled INs (**Fig. 2 b**, see Methods). The number of neurons showing AP firing in response to extrinsic and intrinsic inputs varied between IN populations, with SOM and PV INs activated by both extrinsic and intrinsic inputs and VIP INs almost exclusively activated by extrinsic afferents (**Fig. 2 c-e**). Quantitatively, we found that AP probabilities and the number of APs generated in PCs upon a single stimulation was generally greater following the activation of extrinsic over intrinsic inputs (**Fig. 2 f-g**). On the other hand, SOM INs did not show overall differences in AP probability between extrinsic and intrinsic inputs, although the number of APs evoked by a single intrinsic stimulation was significantly higher relative to extrinsic inputs (**Fig. 2 h-i**). Analogous findings were observed for PV INs (**Fig. 2 j-k**). It is important to note that extrinsic stimulation was still able to trigger APs in both SOM and PV INs but to a smaller degree than local BLA PC stimulation, which in many cases resulted in burst-like firing (see voltage trace in **Fig. 2 b**). In contrast to SOM and PV INs, but similar to PCs, AP probability and number in VIP INs were significantly lower upon intrinsic, relative to extrinsic, stimulation. (**Fig. 2 l-m**). Interestingly, LEC inputs to PCs and VIP INs were less effective at driving APs than other extrinsic inputs. These data confirm that SOM and PV INs are more robustly activated by local BLA PCs than by extrinsic afferents, while VIP INs are almost exclusively activated by extrinsic (non-LEC) inputs and not by local BLA PCs. Furthermore, analysis of evoked AP/EPSP peak latency differences between extrinsic and intrinsic stimulation on INs, relative to the paired PCs, revealed longer peak latencies for extrinsic- than to intrinsic inputs- for SOM, but not for PV or VIP INs (**Fig. 2 n-p** and traces in **a-b**), in accordance with their predominantly disynaptic response to long-range afferents described earlier.

**FIGURE 2.**
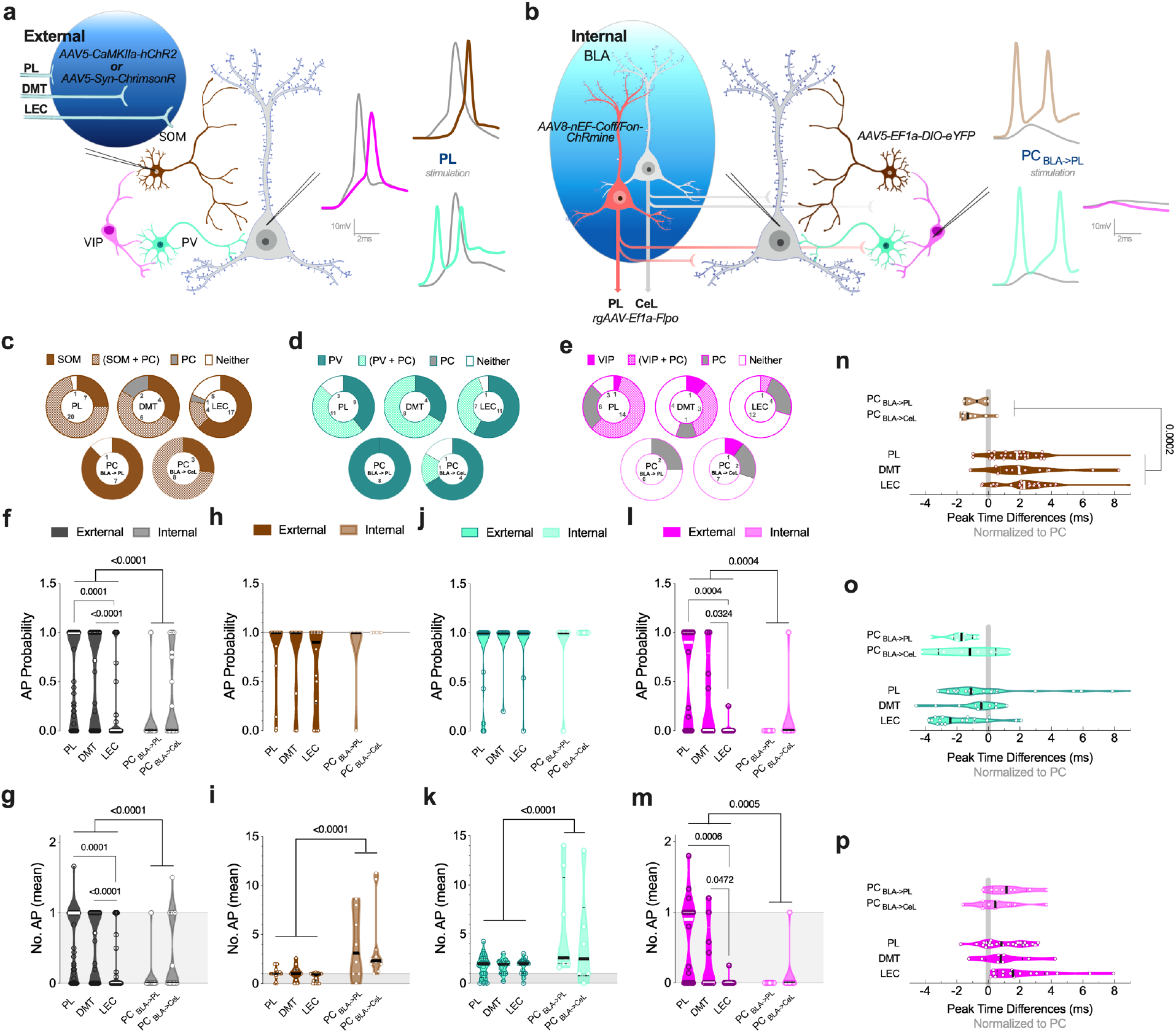
Comparative analysis of internal and external excitatory drive to BLA IN types. a-b, Diagrams of experimental approaches to isolate external and internal afferents to BLA cell types. **c-e,** Proportion of cells exhibiting APs among the recorded pairs in response to extrinsic afferent (PL, DMT, LEC) or local PC (PC _BLA->PL_, PC _BLA->CeL_) stimulation. **f-g,** AP probability and number triggered in PCs in response to extrinsic or intrinsic stimulation. **h-i,** AP probability and number triggered in SOM INs in response to extrinsic or intrinsic stimulation. **j-k,** AP probability and number triggered in PV INs in response to extrinsic or intrinsic stimulation. **l-m,** AP probability and number triggered in VIP INs in response to extrinsic or intrinsic stimulation. **n-p,** Time differences between peak EPSP/AP recorded from SOM (**n**), PV (**o**), and VIP (**p**) INs relative to the peak EPSP/AP of paired PCs upon stimulation of local PC _BLA->PL_ or PC _BLA->CeL_ (top) or extrinsic (PL, DMT, or LEC) inputs (bottom). All data derived from PC-IN pairs (input: n=cell pairs, number of mice): SOM (PL: 21, 12), (DMT: 12, 5), (LEC: 27, 11), (PC _BLA->PL_: 8, 3), (PC _BLA- >CeL_: 11, 4); PV (PL: 23, 11), (DMT: 12, 6), (LEC: 19, 7), (PC _BLA->PL_: 8, 4), (PC _BLA->CeL_: 6, 3); VIP (PL: 24, 8), (DMT: 9, 4), (LEC: 18, 6), (PC _BLA->PL_: 8, 3), (PC _BLA->CeL_: 10, 4). **f-p,** Statistical analysis and P values via two-way ANOVA followed by Sidak post hoc tests (see **Table S1**).

Next, we examined effects of intrinsic BLA stimulation on distinct IN types and neighboring PCs using voltage-clamp recordings and analysis of excitatory and inhibitory postsynaptic current (EPSC/IPSC) amplitudes elicited by stimulation of local PL- or CeL-projecting BLA PCs (**Fig. S2 a**). Isolated EPSCs and IPSCs were recorded from the same neurons at -70mV and +10 mV, respectively, and application of ionotropic glutamate receptor blockers additionally confirmed the disynaptic nature of IPSCs (**Fig. S2 b**). Upon stimulation of either CeL- or PL-projecting BLA PCs, SOM INs showed significantly smaller disynaptic IPSC but comparable EPSC amplitudes, and an overall increase in the excitation/inhibition (E/I) ratio relative to PCs (**Fig. S2 c-d**). PV INs showed both greater EPSC amplitude and smaller disynaptic IPSC amplitude in response to CeL- and PL-projecting PC stimulation and, like SOM INs, exhibited overall higher E/I ratios relative to PCs (**Fig. S2 e-f**). In contrast, although VIP INs also showed significantly lower IPSC amplitude in comparison to BLA PCs, the E/I ratio remained comparable to PCs, likely due to concomitantly lower EPSC amplitudes (**Fig. 2S g-h**). Taken together, these voltage-clamp and current-clamp experiments suggest SOM and PV INs are strongly activated by intrinsic BLA inputs and thus likely participate in lateral or feedback inhibition of BLA PCs, while VIP INs get almost exclusively extrinsic afferents and thus likely participate in feedforward inhibition. All IN types receive significantly less feedback inhibition upon intrinsic BLA PC stimulation compared to adjacent PCs.

Overall, these data indicate that SOM and PV INs are preferentially suited for feedback inhibition. However, their contributions to feedforward inhibition cannot be excluded given both extrinsic and intrinsic inputs can generate APs, albeit to different degrees, in each IN. To explicitly test a role for SOM and PV INs in feedforward inhibition, we recorded from BLA PCs while optically stimulating PL inputs (ChrimsonR, 617nm light) during Cre-dependent kappa-opioid receptor DREADD (KORD)-mediated inhibition of SOM or PV INs in addition to Cre-dependent ChR2 expression to validate the strength of KORD inhibition (**Fig. S3 a**). Bath application of Salvinorin B caused suppression of direct SOM- and PV-mediated (455nm light) GABAergic IPSCs onto PCs as expected, but only inhibited feedforward IPSCs elicited by PL stimulation in PV-Ai14 mice (**Fig. S3 b-c**). Indeed, a direct comparison of the efficacy of Salvinorin B on feedforward inhibition, relative to direct IN to PC inhibition, revealed a significantly greater effect for PV over SOM INs (**Fig. S3 d**). These data further confirm that PV INs participate in feedforward inhibition, while SOM INs do not.

### BLA IN Subtypes Exert Differential Inhibitory Control Over PCs

Our data suggest distinct IN types differentially participate in microcircuit motifs to regulate the coordinated activity of the BLA, however, the degree to which these INs gate extrinsic afferent-induced activation of BLA PCs remains unclear. To address this, we first determined the IN classes providing direct inhibition to BLA PCs via the expression of ChR2 in SOM, PV, or VIP INs and analysis of evoked IPSPs in BLA PCs (**Fig S4 a**). The majority of PC IPSPs were GABA_A_ receptor-dependent (**Fig. S4 b**) and showed voltage-dependence with reversal potentials of -83.5 mV for PV and -96.2 mV for SOM IN activation (**Fig. S4 c-d**). Average IPSP amplitudes recorded from PCs were highest for SOM > PV INs with VIP activation showing minimal responses even at depolarized potentials (**Fig. S4 c, e**). These data indicate that SOM and PV INs inhibit BLA PCs, while VIP generally do not, consistent with previous results ^19^.

We next wanted to examine the gating efficacy and temporal determinants of SOM and PV IN-evoked inhibition on long-range afferent-induced excitation received by BLA PCs. To this end, we expressed ChrimsonR in the PL, DMT, or LEC of SOM- and PV-Ai14 mice, to allow for red light stimulation of long-range afferents to the BLA. Additionally, DIO-ChR2 was injected into the BLA to allow for blue light activation of different IN types (see **Fig. 3 a-c**). Using this approach, we found that simultaneous or near-simultaneous activation of INs in combination with long-range afferents reduced the EPSP amplitudes measured on PCs (see **Fig. 3 d** for example). Upon PL stimulation, SOM and PV IN activation reduced the EPSP amplitudes, with SOM INs exerting greater suppression than PV cells (**Fig. 3 e-f**). Action potential probability was also suppressed by SOM and PV activation following suprathreshold PL stimulation (**Fig. 3 g-h**). Similarly, in response to DMT stimulation both SOM and PV activation was able to reduce the EPSP amplitudes, with SOM INs exerting greater suppression than PV INs (**Fig. 3 i-j**). Action potential probability was likewise suppressed by SOM and PV activation in the presence of suprathreshold DMT stimulation (**Fig. 3 k-l**). In response to LEC stimulation, SOM and PV activation reached the biggest reduction in PC EPSP amplitudes with similar effectiveness. (**Fig. 3 m-n**). Pyramidal cell action potential probability was suppressed by both IN types paired with suprathreshold LEC stimulation (**Fig. 3 o-p**). Additionally, post-hoc analysis revealed significant differences between SOM- and PV-induced suppression of EPSPs occurred at longer latencies, specifically at 100-200ms ahead of DMT and LEC afferent stimulation. This is consistent with our findings showing similar peak times but longer decay times of SOM- relative to PV-evoked IPSPs on PCs (**Fig. S4 f-g**). Control experiments confirmed that dual-opsin-based approaches were not contaminated by light cross-over activation at light intensities used, and that in the absence of ChR2 expression in INs, ChrismonR-triggered afferent EPSPs were not suppressed by blue light stimulation (**Fig. S4 h-l**). These data indicate that SOM and PV INs can inhibit BLA PC activity and gate sub- and suprathreshold activation of BLA PCs from long-range afferents over long time scales.

**FIGURE 3.**
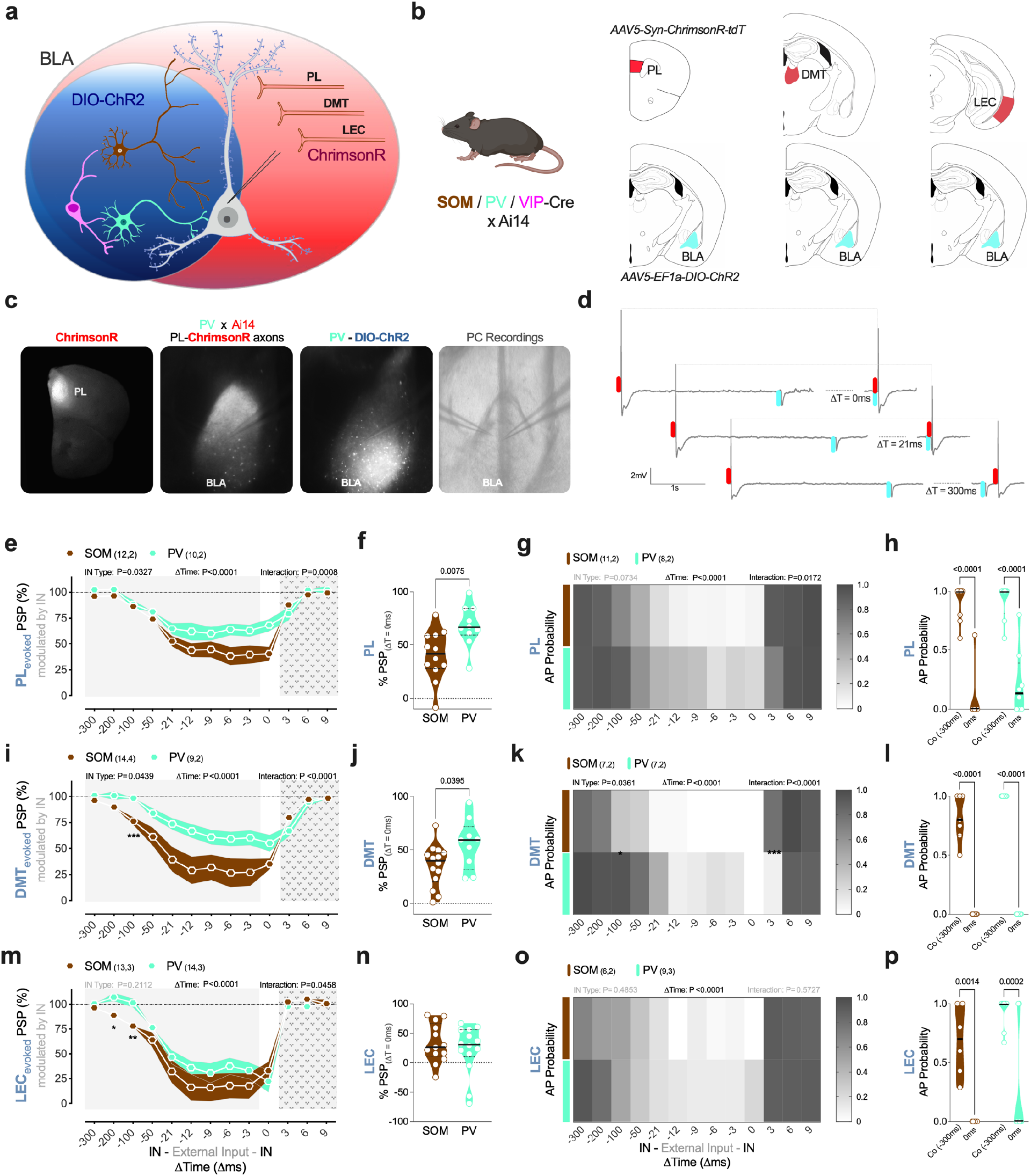
Suppression of extrinsic afferent excitation-induced activation of BLA PCs by different IN types. a-b, Diagram of experimental design. The excitatory opsin ChrimsonR encoding AAV was injected into the PL, DMT, or LEC and a Cre-dependent ChR2 was injected into the BLA of SOM-, PV- or VIPxAi14 mice. Whole-cell current-clamp recordings were obtained from BLA PCs. **c,** Representative photomicrographs of PL-ChrimsonR injection site, ChrimsonR terminals and PV cells, PV-ChR2 expression, and DIC image of BLA PC recordings. **d,** Representative traces (related to **c**) of dual-opsin experiments showing extrinsic PL afferent- induced EPSPs (red-light), PV activation-induced IPSPs (blue-light), and mixed responses to blue-and red-light combinations given at different time intervals. **e-f,** Effects of SOM and PV IN activation on PL-triggered PSPs as a function of blue-red light stimulation interval relative to PL stimulation, and maximal PSP inhibition at ΔT=0ms for SOM and PV INs. **g-h,** Effects of SOM and PV IN activation on PL-triggered AP probability as a function of blue-red light stimulation interval relative to PL stimulation, and maximal AP probability differences between ΔT=300ms (control; Co) and ΔT=0ms for SOM and PV INs. **i-l,** same as **e-h** for DMT-induced PSP and APs in BLA PCs. **m-p,** Same as **e-h** for LEC-induced PSP and APs in BLA PCs. Sample sizes (n cells, N mice); **e-f,** SOM (12,2), PV (10,2); **g-h,** SOM (11,2), PV (8,2); **i-j,** SOM (14,4), PV (9,2); **k-l,** SOM (11,2), PV (7,2); **m-n,** SOM (13,3), PV (14,3); **o-p**, SOM (6,2), PV (9,3). Statistical analysis **e, g, i, k, m, o** via two-way ANOVA followed by Sidak post hoc test comparing IN types; **f, j, n,** via unpaired t-test; **h, l, p** via two-way ANOVA followed by Sidak post hoc test comparing AP probabilities under control (Co) and ΔT=0ms conditions (see **Table S1**).

### BLA IN Subtypes Exhibit Unique Activity Patterns Across Auditory Cue Fear Conditioning

Given the substantial differences observed in extrinsic and intrinsic connectivity between specific BLA IN types as well as the diverse inhibitory effects of these INs onto PCs, we reasoned that different IN classes would display distinct activity changes during associative learning^13–16^. Mice were injected with Cre-dependent GCaMP7f encoding AAV followed by optical fiber implantation into the BLA to measure the activity of SOM, PV or VIP INs during an auditory cue fear conditioning paradigm (**Fig. 4 a-b and Fig. S5 a**). During conditioning, SOM INs showed progressive increases in both CS+- and US-evoked responses across conditioning (**Fig. 4 c-d**). Comparisons of peak z-score and AUC for onset (0-5s) and duration (5-29s) of CS+ presentations showed progressive increases in activity between the first and last CS-US pairing, with no signals detectable following the first tone (**Fig. 4 e**). US-evoked responses exhibited a similar sensitizing pattern with higher persistent activity, strikingly lasting up to 30s, following the brief 1s shock (**Fig. 4 c-d, f-g**). In contrast to SOM neurons, PV INs displayed consistent CS+-evoked signals across conditioning trials (**Fig. 4 h-j**). PV US responses, on the contrary, showed a significant reduction in peak z-score and AUC from early to late presentations (**Fig. 4 k**), mostly limited to the 5s following shock onset (**Fig. 4 l**). VIP INs showed minimal CS+-related activity (**Fig. 4 m-o**), but their robust US response followed a similar pattern as PV INs: decaying phasic activity across conditioning trials (**Fig. 4 p-q**). Averaged data across all conditioning trials from individual male and female mice support the reliability of responses across subjects (**Fig. S5 b-e**). YFP-expressing control mice showed no overall CS+- or US-evoked activity across conditioning (**Fig. S5 f-l**). These data indicate differential fear-conditioning-associated changes in IN activity across learning, with SOM INs displaying progressive and long-lasting increases in CS+- and US-evoked activity across conditioning, and VIP INs showing primarily sharp, short-lived, US-evoked signals that habituate across CS-US pairings. Interestingly, PV INs exhibit some features of both VIP (sharp habituating US response) and SOM (late CS+ response) INs.

**FIGURE 4.**
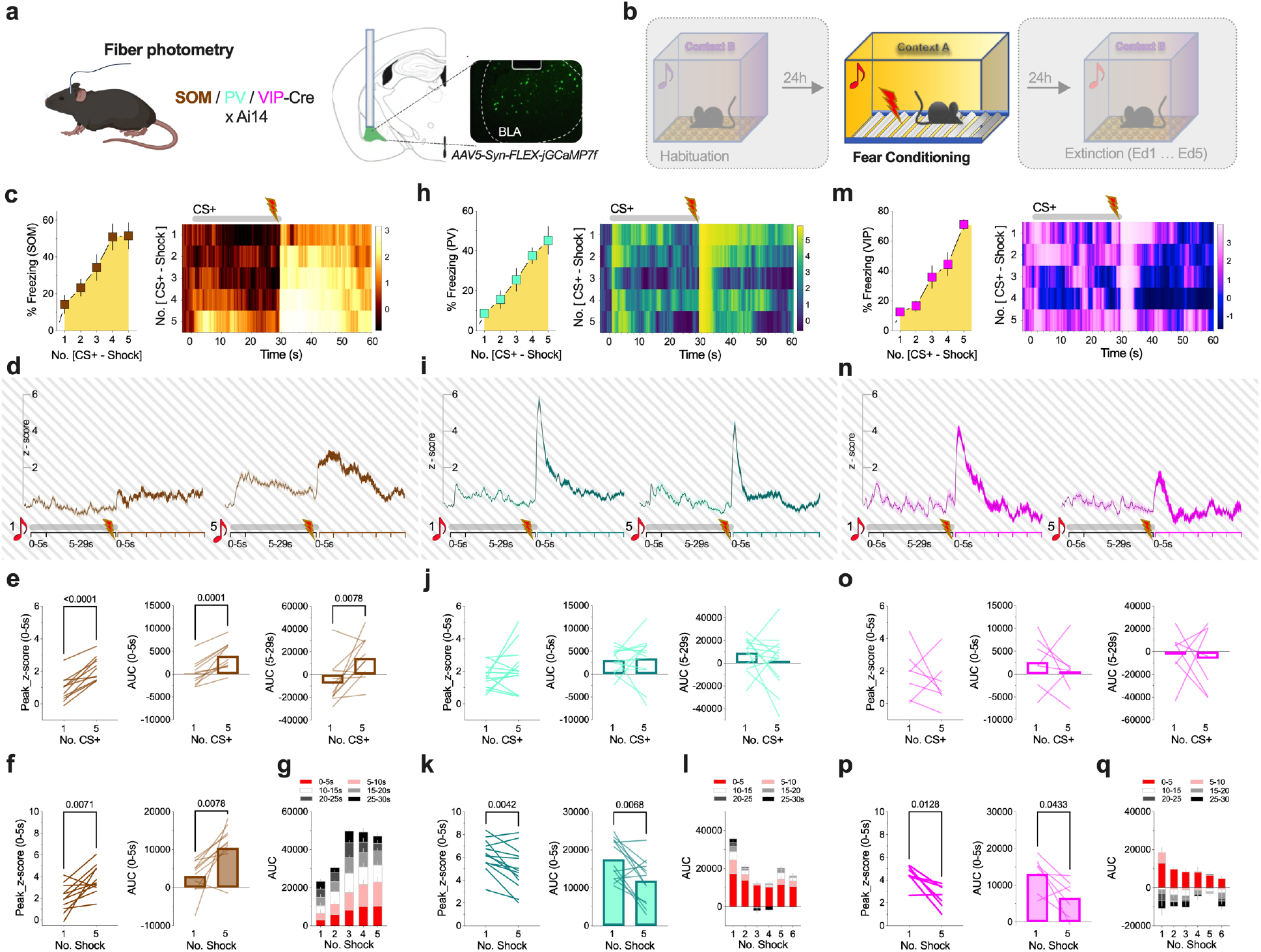
Activity of distinct IN types across fear conditioning. a-b, Schematic diagram of experimental design. In vivo calcium levels were monitored via Cre-dependent expression of GCaMP7f using fiber photometry recordings of SOM, PV and VIP BLA INs during fear conditioning. **c,** Freezing data and heatmap of z-score values across 5 CS-US pairings for SOM INs. **d,** Average z-score traces of SOM activity during CS-US pairing number 1 (left) and 5 (right). Data was averaged across mice per the number of CS+-shock presentation, z-score traces represent mean ± SEM. **e,** Peak z-score and AUC early (0-5s) and late (5-29s) responses to CS+ No. 1 and 5. **f,** Peak z-score and AUC responses to US (shock) No. 1 and 5. **g,** AUC distribution of neuronal activity following shock presentations over 30s averaged for every 5s in SOM INs. **h-l,** same as **c-g** for PV INs. **m-q,** same as **c-g** for VIP INs. Sample size (mice; sex); SOM (n=11; 6M/5F), PV (n=13; 7M/6F), VIP (n=7; 3M/4F). Statistical analysis and P values via two-tailed paired t-test throughout.

### BLA IN Subtypes Exhibit Unique Activity Patterns Across Conditioned Fear Extinction

During extinction training, another set of unique activity patterns emerged among the IN types (**Fig 5 a-b**). SOM INs responded with sharp, phasic (0-3s) increases in activity following CS+ presentations throughout extinction days (early day 1 vs. late day 5) (**Fig. 5 c-d**). In contrast, a slower, long-lasting reduction in calcium signals was detected over the second half of CS+ presentation (15-30s) during early phase of extinction, which attenuated with extinction learning (**Fig. 5 c-d**). Additionally, upon CS+ offset, another sharp phasic response was detected in SOM INs during the early, but not late, phase of extinction (**Fig. 5 c** inset and **e**). Interestingly, the AUC analysis of slow reductions (15-30s) showed indirect correlations with the overall CS+-induced freezing values, with larger reductions at higher freezing levels, and smaller reductions at lower freezing levels. (**Fig. 5 f**). The phasic CS+ offset-dependent increases, on the other hand, indicated direct correlations with the tone-induced freezing and completely attenuated with extinction training, when the freezing lessened (**Fig. 5 g**). Direct comparisons of CS+ onset and offset peaks revealed similar magnitude events during early, but not late, extinction (**Fig. S6 a**); notably, the CS+ offset peak coincided with increased movement after tone termination (**Fig. S6 a** inset). PV INs exhibited similar activity patterns to SOM neurons on early extinction day for both the phasic CS+ onset rise and the late long-lasting decay in calcium signals during tone presentations (**Fig. 5 h**). Both, however, attenuated by late extinction trials (**Fig. 5 i**). Like in SOM INs, late CS+ decline in PV IN activity indirectly correlated with freezing levels, signifying increased PV neuronal activity with the development of extinction learning compared to fear state (**Fig. 5 k**). CS+ offset response, although small, revealed a direct correlation with freezing, both decreased by late extinction (**Fig. 5 l** and **Fig. S6 b**) Analysis of VIP INs demonstrated solely a phasic increase upon CS+ presentation that weakened with the development of extinction (**Fig. 5 m-n**), and CS+ offset yielded to no changes in calcium activity in these cells (**Fig. 5 o** and **Fig. S5 c**). CS+ onset peak showed direct correlation with freezing level, both attenuated with extinction (**Fig. 5 p**), and lack of late-onset CS+ shift (15-30s) or CS+ offset response meant no correlations with freezing for VIP INs (**Fig. 5 q**). These data indicate differential and dynamic changes to CS+ presentation across extinction training in SOM, PV, and VIP INs. Analogous to the distinction in activity patterns during fear conditioning, SOM and VIP INs displayed markedly distinct activity patterns and correlations with freezing levels, whereas PV INs showed an intermediate phenotype with properties of both SOM (late CS+ slow reductions) and VIP INs (habituating CS+ onset response). No consistent changes in activity related to CS+ onset or offset were observed in YFP control mice across extinction (**Fig. S6 d-f**).

**FIGURE 5.**
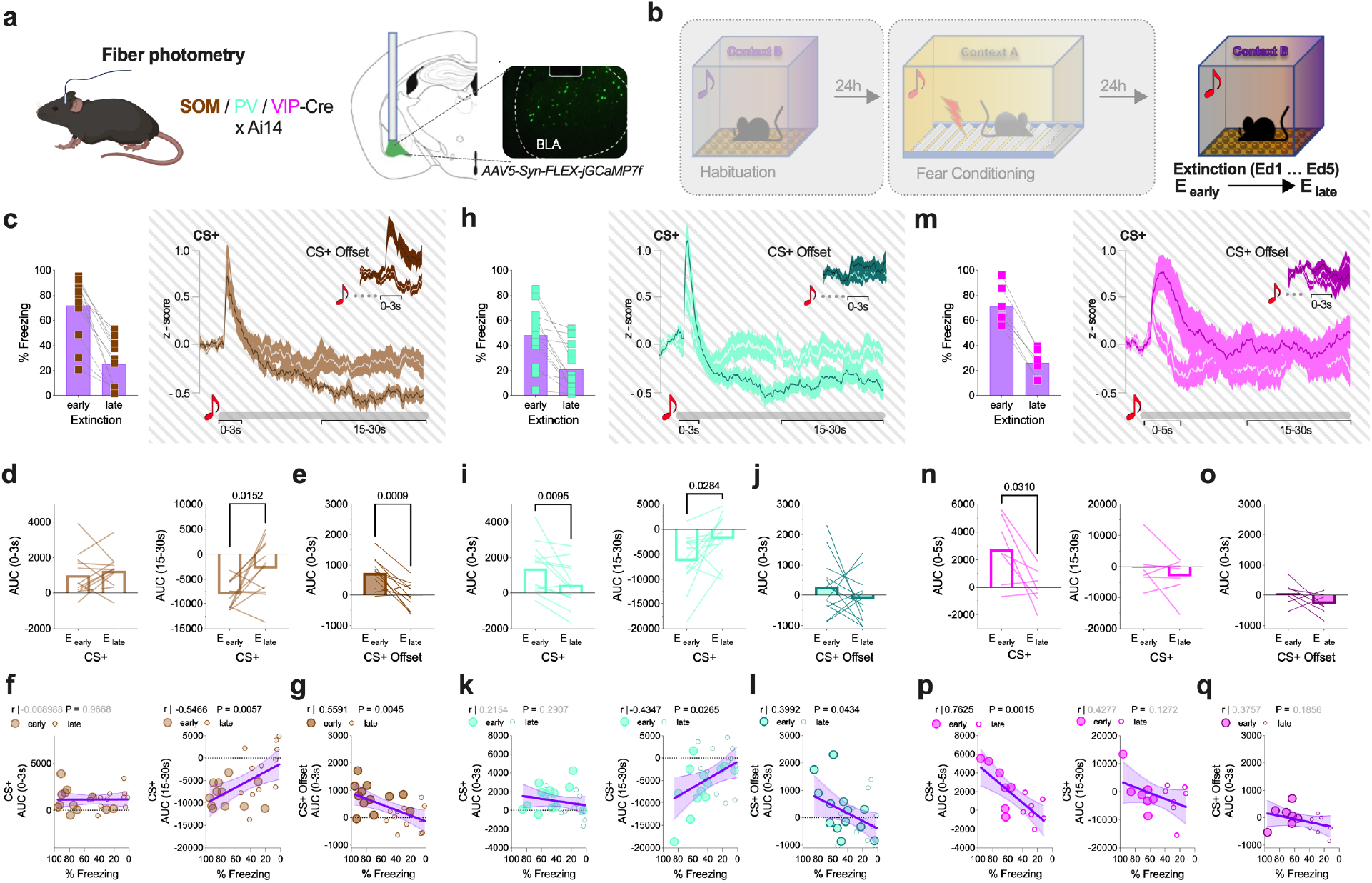
Activity of distinct IN types across fear extinction. a-b, Schematic diagram of experimental design. In vivo calcium levels were monitored via Cre-dependent GCaMP7f expression using fiber photometry recordings of SOM, PV and VIP INs during fear extinction (4-5 days). **c,** Freezing reductions across extinction and average z-score traces of SOM IN activity during early (day 1; dark line) and late (d5, white line) extinction days upon CS+ onset and offset (insert). Z-score traces of CS+ onset and offset were first averaged per session, then across mice, traces represent mean ± SEM. **d,** AUC following CS+ presentation (0-3s and 15-30s). **e,** AUC (0-3s) for CS+ offset. **f,** Linear regression for CS+ AUC (0-3s) vs. freezing, CS+ AUC (15-30s) vs. freezing, and **g,** CS+ offset AUC (0-3s) vs. freezing. **h-l,** same as **c-g** for PV INs. **m-q,** same as **c-g** for VIP INs. Sample size (mice; sex); SOM (n=11; 6M/5F), PV (n=13; 7M/6F), VIP (n=7; 3M/4F). Statistical analysis via two-tailed paired t-test (**d-e**, **i-j**, **n-o**) and via Pearson r correlation and simple linear regression (**f-g**, **k-l**, **p-q**).

### BLA IN Subtypes Show Differential Responsivity to Behavioral State Transitions

While our analysis above demonstrates discrete changes in calcium activity accompanying CS+ presentation (onset, during and offset) and reveals correlations to overall CS+ freezing levels, the close relationship between IN activity and freezing behavior remained unclear in that evaluation, as mice exhibit multiple transitions between freezing and non-freezing states during the 30-second CS+ presentations and freezing in between tones (inter-tone-intervals). To address the question whether IN activity is directly related to behavioral transitions in addition to sensory cue presentations we measured SOM, PV, and VIP IN activity specifically during freezing and moving states, as well as at the transitions from moving to freezing and freezing to moving states across our fear conditioning and extinction protocol (**Fig. 6 a**). Our analysis excluded the first few seconds of CS+-, shock onset and offset periods to reduce confounding from sensory signals (see above) and distinguished between tone and inter-tone-intervals.

**FIGURE 6.**
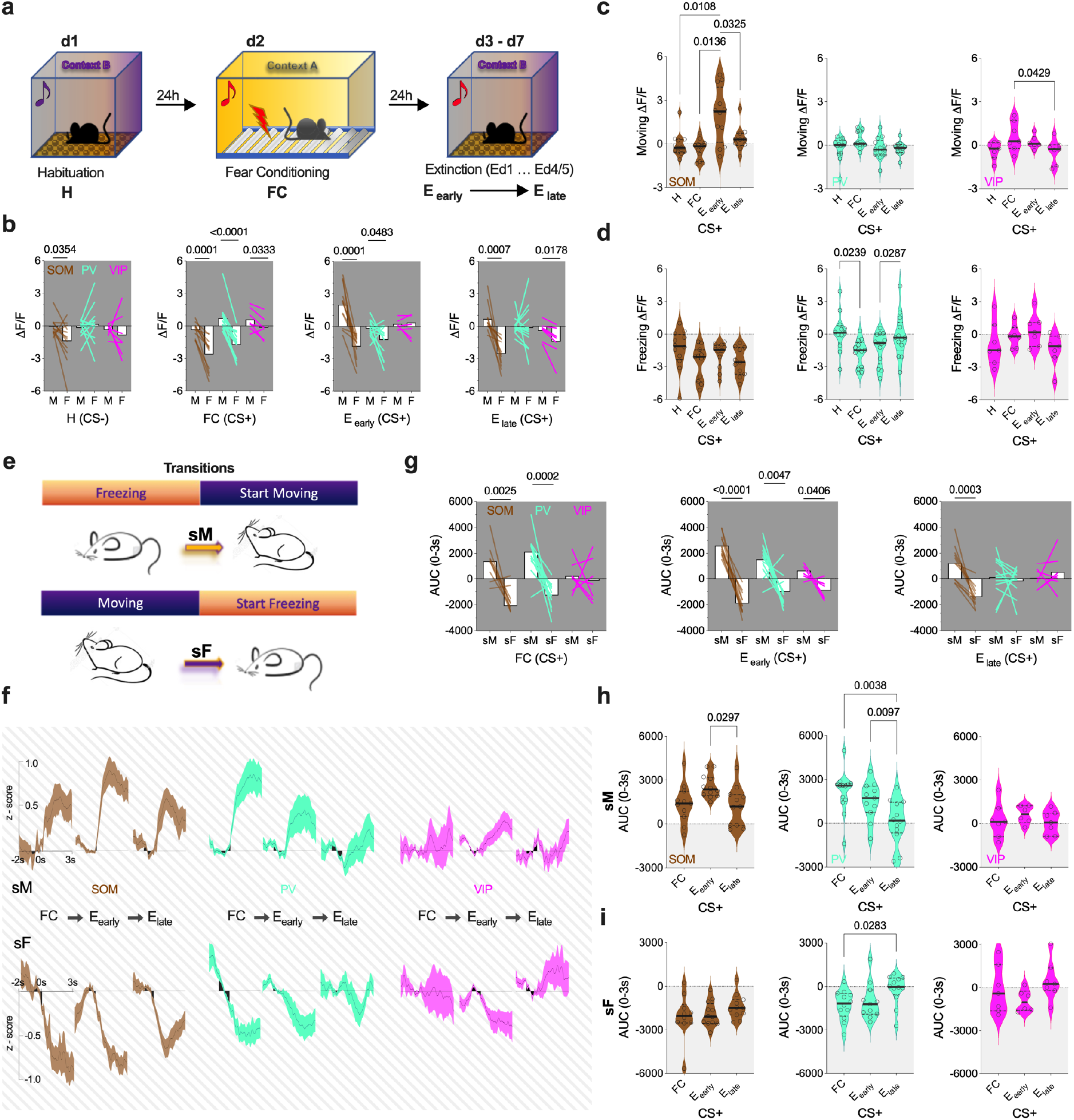
Behavioral state transition-related activity of distinct IN types during conditioned stimulus presentation. a, Diagram of task days during which behavioral state transition data was analyzed. **b,** Averaged ΔF/F values during bouts of motion (M) and freezing (F) on habituation (H), fear conditioning (FC), early (E_early_) and late extinction (E_late_) days recorded from SOM (brown), PV (green) and VIP (pink) INs. **c,** Averaged ΔF/F values during bouts of motion during CS presentations compared across behavioral days for SOM, PV and VIP populations. **d,** Averaged ΔF/F values during bouts of freezing compared across behavioral days for SOM, PV and VIP populations. **e,** Diagram of behavioral state-transition analysis depicting dissection of transitions where mice start moving (sM) or start freezing (sF). Data analysis for habituation day was omitted due to low number of transitions per mouse. **f,** Averaged z-score traces during behavioral state transitions for SOM, PV and VIP populations on fear conditioning, early and late extinction days. Z-score traces of transitions were first averaged per session, then across mice; traces represent mean ± SEM. Data were baselined to 1 second prior to behavioral state transition as shown in black. **g,** AUC (0-3s) values of start moving and start freezing transitions during CS+ presentations compared within IN types on separate days for SOM, PV and VIP populations. **h,** AUC (0-3s) values following motion onset (sM) during CS+ presentations compared across days for SOM, PV and VIP INs. **i,** AUC (0-3s) values following freezing onset (sF) during CS+ presentations compared across days for SOM, PV and VIP INs. Sample size (n= mice; sex); SOM (n=11; 6M/5F), PV (n=13; 7M/6F), VIP (n=7; 3M/4F). **b, g,** P values and analysis via paired t-test comparing M to F and sM to sF for each IN type; **c, d, h, i,** analysis via one-way ANOVA followed by Tukey post hoc test comparison across task days (see **Table S1**).

We started by quantifying the calcium signal dynamics of distinct IN populations as a function of motion and freezing during habituation (H), fear conditioning (FC), and extinction days (E_early_ and E_late_) during CS+ presentation. The additional inclusion of habituation day dataset allowed us to assess data from full range of fear states (low-high-low). ΔF/F values were averaged during motion and freezing behaviors across sessions and compared within mice. Analysis revealed a divergent pattern in calcium signals with generally higher ΔF/F values during movement and lower values during freezing, depending on the day of behavioral task and cell type. Specifically, on habituation day, only SOM INs exhibited lower ΔF/F values during freezing relative to motion throughout CS- presentations (an auditory stimulus only present in the contexts of H and E days) (**Fig. 6 b**). On the fear conditioning day, remarkably, each IN type revealed a split between overall calcium signals derived from freezing and motion bouts during CS+ presentations, albeit small for VIP, the split remained notable in SOM and PV INs on the following day (E_early_), when mice expressed the highest level of fear (**Fig. 6 b**). After the development of fear extinction (E_late_) SOM INs (and to a lesser degree VIP INs) still showed attenuated signals during freezing relative to motion (**Fig. 6 b**). Direct comparisons of movement-derived ΔF/F values across the days within IN classes revealed increased SOM IN activity on the day of highest fear expression (E_early_), relative to other task days, while VIP IN activity indicated slight separation between fear conditioning (higher) and late extinction day (lower) (**Fig. 6 c**). Freezing-associated SOM IN calcium signals showed comparable attenuations across days, whereas PV INs showed attenuations on days associated with high fear expression (FC and E_early_) relative to days with low fear expression (H and E_late_). VIP IN calcium signals displayed no distinctions across sessions (**Fig. 6 d**).

To explore whether these patterns emerged from discrete time-locked events, we examined the transitions between movement and freezing behaviors (**Fig. 6 e**). We analyzed events with a minimum of 1s stable behavioral states both before and after transitions in order to evaluate well-defined transitions and due to minimal freezing expression, the habituation day dataset could not be assessed. Indeed, the analysis revealed a sharp increase in calcium activity accompanying the freezing to movement transitions, whereas a sharp decrease coincided with the movement to freezing transitions. These phenotypes were most notable in SOM INs followed by PV INs, with minimal distinctions in VIP cells (**Fig. 6 f)**. AUC (0-3s) values of z-scored calcium traces were used for quantification at behavioral transitions and averaged across sessions per mouse during or between CS+ presentations. AUC values showed marked separations between transition types (start moving/start freezing; sM/sF) on each day for SOM neurons, on fear conditioning and fear recall day (E_early_) for PV cells, and only on fear recall (E_early_) for VIP cells (**Fig. 6g**). Direct comparisons of freezing to moving transitions across days within IN classes revealed larger increase in SOM neuronal activity on early vs. late extinction day, higher activities on both fear conditioning and early extinction vs. late extinction for PV cells, and no significant changes in VIP neurons (**Fig. 6 h**). Comparisons of moving to freezing transitions showed separation only between fear conditioning and extinguished fear association (E_late_) days for PV cells (**Fig. 6 i**). Similar quantitative effects were observed in all measures during inter-tone intervals (**Fig. S7**). YFP-expressing control mice displayed no changes in activity between times of motion and freezing (**Fig. S7 a**) with small inconsistent changes at transition points (**Fig. S7 f**). Taken together, these data illustrate both SOM and PV INs undergo marked behavioral state transition-related activity changes with sharp increase and decrease at the start of movement and freezing, respectively. Interestingly, these signals were present on each day in SOM neurons, but only emerged on days with heightened fear expression in PV neurons (and to a much lesser extent in VIP neurons), indicating a dynamic activity pattern tracking internal state. Additionally, these behavioral state-associated activity changes were independent of CS+ presentation, underscoring their independence from sensory-driven activity.

## DISCUSSION

While the BLA is a well-established hub critical for the acquisition and extinction of conditioned fear responses^1,3,5,7^, a comprehensive analysis of how specific BLA GABAergic INs contribute to different circuit motifs and the relationship of these motifs to the functional properties of IN classes across conditioned fear acquisition and extinction learning is lacking. Moreover, learning associated plasticity within the IN classes and their potential differential contributions to the representation of conditioned freezing and related behavioral state transitions has not been explored. Here, we address these open questions via investigation of functional synaptic connectivity and microcircuit organization of BLA SOM, PV, and VIP INs, and their responses to sensory cues and behavioral state transitions across fear acquisition and extinction.

Our electrophysiological analysis of IN synaptic connectivity and microcircuit organization support a model whereby SOM INs prominently participate in feedback inhibition of BLA PCs, VIP INs contribute primarily to feedforward disinhibition of PCs, and PV INs engage in both feedforward and feedback inhibition in the BLA (**Fig. S8**). This model is supported by multiple pieces of convergent data. First, we observed small direct monosynaptic, yet robust and consistent disynaptic excitation of SOM INs upon stimulation of all extrinsic inputs tested, whereas VIP INs received almost exclusively monosynaptic long-range excitation; and PV INs displayed both strong monosynaptic and disynaptic excitation depending on the afferent input. Second, direct analysis of the probability and number of APs generated in response to intrinsic versus extrinsic stimulation shows that, while both intrinsic and extrinsic sources can readily trigger APs in SOM and PV neurons, activation of local PCs evokes a higher number of APs in both IN types. VIP INs could not be triggered by intrinsic afferents, and their firing probabilities showed a similar profile to those of PCs. Third, voltage-clamp studies revealed that both SOM and PV INs exhibit high E/I upon stimulation of local intrinsic BLA PCs, but VIP INs do not. Fourth, unlike VIP INs, SOM and PV cells provide effective and long-lasting inhibition of PCs. Finally, our chemogenetic data imply that PV, not SOM, INs participate in feedforward inhibition of BLA principal cells driven by extrinsic afferents.

This model aligns with several previous findings. Monosynaptic rabies tracing suggests robust local innervations of SOM INs ^19^, supporting our data that SOM INs preferentially participate in feedback inhibition of BLA PCs, similar to their role in the neocortex.^24,25^. With regard to BLA PV INs, research in cortical areas reinforce the roles for PV INs in both feedback and feedforward inhibition ^24,25^, and anatomical and functional BLA studies have revealed strong excitatory inputs from local PCs on PV INs ^26–29^, supporting the mixed phenotype we observed depending on the examined terminals. Anatomical data have also revealed BLA VIP INs receiving VGluT2 (likely of thalamic origin) and VGluT1 cortical synaptic contacts^30^, consistent with our electrophysiology data showing prominent monosynaptic excitation by extrinsic afferents only. Finally, previous studies corroborate our findings that while SOM and PV INs can provide powerful inhibition to BLA PCs ^19,27^, VIP INs generally do not ^19,30^. This suggests that VIP INs are primarily involved in feedforward inhibition onto other INs ^19^, mediating feedforward disinhibition of PCs, as similarly described in other cortical-like structures ^25^.

In partial contrast to our results, a recent study suggested that a fast-spiking subtype of SOM, but not PV, INs mediate feedforward inhibition onto BLA PCs upon LEC stimulation ^28^. We found that, although SOM INs could be activated by LEC inputs, disynaptic responses were far more robust. These discrepancies could be due to the methodological differences. For example, our data derive from optical stimulation of the fibers originating from the mid-posterior LEC, while the earlier study utilized electrical stimulation in horizontal slices. Specific recording sites within the basolateral complex could also account for these differing results. In support of this contention, another study using electrical stimulation in coronal sections found divergent roles for PV INs in feedforward inhibition in the lateral relative to basolateral nucleus ^31^. This is consistent with our finding that PV INs show varying anatomical substrates for feedforward inhibition depending on the afferent source. Furthermore, a recent study reported PL-induced increase in BLA SOM IN activity contributing to the discrimination of non-threatening stimuli ^18^. PL-induced activation of SOM INs is apparent in our data, although the strongest activation results from disynaptic excitation.

To gain insight into how the synaptic organizational motifs of specific IN types may relate to their functional properties, we utilized fiber photometry-based calcium measurements from SOM, PV, and VIP INs across a conditioned fear acquisition and extinction learning task. During fear conditioning, CS+ calcium responses showed sensitization in SOM INs, remained stable in PV, and were minimal in VIP INs. Upon US deliveries, responses again showed sensitization in SOM INs but displayed habituation in both PV and VIP INs. Interestingly, the CS+- and US-evoked calcium activity in SOM population lasted substantially longer than in PV or VIP neurons. These data highlight divergent response and plasticity profiles of specific IN populations, with SOM and VIP cells displaying the most distinct phenotypes and PV INs contributing to a mixed representation of SOM and VIP INs, namely CS+ responsivity and habituative US responses, respectively. These data are consistent with previous reports that habituating US responses in VIP cells may represent attenuating US salience as a function of predictability ^19^, and studies demonstrating dynamic activity patterns of subpopulations of BLA SOM and PV INs in response to CS+ and US presentations ^17,19^. Lastly, sensitizing increases in SOM IN activity across fear conditioning have also been reported in the prefrontal cortex ^32^, raising the intriguing possibility that these learning associated changes may represent a uniform property of SOM INs across brain regions.

After conditioning, mice underwent extended extinction training which resulted in reduced freezing in all mice accompanied by distinct changes in activity of different IN types. In the early phase of extinction, when freezing levels were high, we discerned three types of activity patterns associated with CS+ presentation. First, a phasic spike upon CS+ onset which was observed in all three IN types. Second, a subsequent slow reduction in activity for the duration of the tone found in SOM and PV INs. Third, a CS+ offset spike, present most prominently in SOM INs. Interestingly, dynamic changes in these activity patterns were observed between early and late extinction trials. The first, phasic CS+ onset spike, remained stable in SOM but habituated in PV and VIP INs. The second, decreasing signal during CS+ presentation, went through habituation in both SOM and PV INs. The third, CS+ offset peak, became absent in SOM neurons by late phase of extinction. Yet again, SOM and VIP INs displayed the most distinct dynamic activity patterns with PV INs exhibiting a mixed profile of both SOM INs (slow CS+-related reduction in activity that habituated across extinction) and VIP INs (sharp CS+ phasic signal that habituated across extinction). In addition, examination of the relationships between these calcium activity patterns and overall freezing levels revealed a direct correlation upon CS+ onset in VIP INs and a negative correlation during tones in SOM and PV INs, when their calcium signals displayed varying degrees of reduction. In other words, VIP IN activity decreased while SOM and PV activity increased (i.e. less reduction in calcium signal) with the development of extinction learning. Furthermore, a direct correlation was observed at CS+ offset peak in both SOM and PV INs between freezing level and calcium activity.

Next, we explored the activity of specific IN types as a function of freezing and moving, and during transitions between these states. We started out by assessing the overall activity during times of motion and freezing and, depending on the trial and cell type, found a distinct separation in activity levels, with higher signals during movement and lower signals during freezing. Notably, in SOM neurons this segregation was present across all days, even during habituation when freezing levels were minimal. The degree of divergence between moving/freezing calcium signals, however, markedly increased with the expression of fear, during fear conditioning and early extinction, with a similar separation also present in PV INs. VIP INs, on the other hand, showed small changes that did not systematically vary with the behavioral training. To explore whether these activity patterns originated from discrete time-locked events, we examined the transitions between movement and freezing behaviors. Indeed, sharp increase accompanied the freezing to movement transitions, and sharp decrease coincided with the movement to freezing transitions on each day in SOM INs, during fear conditioning and early extinction in PV INs, and during early extinction in VIP INs. These responses were present not only during, but in between tone periods, indicating they are not related to sensory input. Interestingly, a recent study reported congruent mobility-related activity changes of these INs in tail suspension test, and other work has demonstrated increased SOM IN activity during presentation of safety predicting cues associated with suppressed freezing ^18,33^. Moreover, chemogenetic inhibition of BLA SOM INs increases conditioned freezing behavior ^34^, in line with our data revealing low SOM IN activity during freezing.

Various functional implications of our data are worth mentioning. First, studies have highlighted the presence of valence-coding ensembles of BLA PCs likely utilizing subpopulations of GABAergic INs to generate lateral inhibition and support functional antagonism between such ensembles ^12,35^. Given that SOM and PV INs are strongly activated by local PCs they could contribute to valence coding through lateral inhibition if they preferentially target opposing valence ensembles. Determining whether these INs are anatomically positioned to promote functional antagonism of opposing ensembles require future investigations. Second, that intrinsic inputs can evoke multiple APs in SOM and PV INs, combined with their ability to markedly modulate the effectiveness of extrinsic afferent-induced activation of BLA PCs (up to 200ms), highlights the ability of these INs to dramatically constrain BLA PC activity and potentially act as low pass filters in the face of robust extrinsic excitation. Third, we show that SOM and PV INs respond to both sensory cues as well as behavioral state transitions suggesting multidimensional coding properties for subtypes of BLA INs, as has been previously reported for BLA PCs ^36,37^, however adequate testing of this hypothesis will require detailed single-cell level analysis of IN activity. Fourth, while we detected clear fluctuations in IN activity as a function of behavioral state (i.e. freezing or moving), the magnitude of these changes was dynamic across conditioning and extinction for some cell type. One interpretation of these data is that the magnitude of these transitions reflects an internal state not captured by a binary behavioral classification, consistent with recent studies demonstrating state-dependent coding within BLA PCs ^38,39^. In other words, if alterations in IN activity reflect only or purely behavioral transitions, the magnitude of these changes would not be expected to vary across conditioning and subsequent extinction as fear states decline. Lastly, our data demonstrate that SOM IN activity (and to a lesser degree PV activity) tightly reflects freezing onset and offset with decreases and increases, respectively. Similarly, increases in SOM IN activity have been reported during learned safety cues causing desynchronization of BLA PC firing, and consequently, blocking theta oscillation phase reset^18^. Thus, it is plausible that suppression of theta power could correspond to increased motion and SOM activity, while enhancement of theta power would yield decreased motion and SOM activity. Further studies are necessary to dissect the causal relationships between IN activity, synchronization, and behavior across fear acquisition and extinction.

In summary, our data reveal a synaptic/circuit organization-function relationship diversity among BLA SOM, PV, and VIP INs. Specifically, we show that SOM INs mediate feedback inhibition and exhibit the greatest degree of learning-induced plasticity changes displaying multiplexed representations of both salient sensory stimuli and behavioral state transitions that are dynamic across acquisition and extinction of conditioned fear. VIP INs, that mediate feedforward disinhibition, respond primarily to salient sensory cues, while PV INs participate in both feedback and feedforward inhibition and exhibit functional properties of both SOM and VIP INs throughout our behavioral assay. Considering the subpopulations of PV cells reported ^40–42^, it is possible that distinct subpopulations contribute to the mixed functional phenotypes of PV INs observed herein. For example, PV INs mediating feedback inhibition could exhibit in vivo activity patterns more akin to SOM INs (e.g. responses to behavioral state transitions, and CS+-related slow reductions during fear recall), while PV cells conveying feedforward inhibition could appear much like the activity of VIP INs (e.g. habituating US responses, and lack of freezing onset/offset-associated changes). Additionally, whether the examined microcircuit organization-function relationships persist across other brain regions or other forms of learning remains to be assessed. Altogether, our findings provide insight into the relationships between distinct IN microcircuit connectivity motifs and in vivo activity patterns across aversive associative learning which could serve as a platform for further understanding synaptic and circuit-level mechanisms underlying stress and trauma-related disorders.

## METHODS AND MATERIALS

### Animals

All experiments were approved either by the Vanderbilt University Institutional Animal Care and Use Committees or by the Northwestern University Animal Care and Use Committee and were conducted in accordance with the National Institute of Health guidelines for the Care and Use of Laboratory Animals. Adult (10+ weeks) male mice were used for electrophysiology experiments and were housed on a 12:12 light/dark cycle (Vanderbilt University) and adult mice of both sexes were used for fiber photometry behavioral experiments housed on a 14:10h light/dark cycle (Northwestern University). Sex differences were not a primary analysis variable in this study and no overt sex differences were observed, thus all data were pooled from both sexes. All mice were group housed by sex (2-5 mice per cage) in a temperature- and humidity-controlled environment with ad libitum access to food and water; and all experiments were performed during the light cycle.

All homozygous transgenic lines were purchased from The Jackson Laboratory. SOM-IRES-Cre (JAX stock #013044), PV-IRES-Cre (JAX stock #017320) and VIP-IRES-Cre (JAX stock #010908) mice were crossed in-house with Ai14 (JAX stock #007914) to obtain SOM:Ai14, PV:Ai14 and VIP:Ai14, respectively. This allowed for Cre-driven expression of the red fluorophore, tdTomato, in the specific IN types. Part of the local excitation dataset (Fig. 2 and Fig. S2) was acquired from SOM-IRES-FLPo heterozygous mice (JAX stock #031629 crossed in-house with wild-type C57BL/6J). For the sake of simplicity, the schematic figures contain the viral strategies used in all three Ai14 crossed IN-Cre lines. See the SOM-FLPo viral strategy in the section of Viruses below. SOM data from these two sources were pooled.

### Viral vectors

The following viruses were gifts from Karl Deisseroth: AAV5-CaMKIIa-hChR2(H134R)-EYFP (Addgene plasmid #26969); AAV5-EF1a-double floxed-hChR2(H134R)-EYFP-WPRE-HGHpA (#20298); AAV5-Ef1a-DIO EYFP (#27056); rgAAV-EF1a-mCherry-IRES-Flpo (#55634) and rgAAV-EF1a-Flpo (#55637) were used combined. AAV8-nEF-Coff/Fon-ChRmine-oScarlet was a gift from Karl Deisseroth & INTERSECT 2.0 Project (#137160); AAV5-Syn-ChrimsonR-tdT was a gift from Edward Boyden (#59171); AAV8-hSyn-dF-HA-KORD-IRES-mCitrine (#65417) was gift from Bryan Roth. In SOM-FLPo heterozygous mice the following viral combination was used for local pyramidal cell excitation in addition to SOM:Ai14 Fig. 2 and Fig S2 depiction: rgAAV-hSyn-Cre-P2A-dTomato (#107738; in PL or CeL), a gift from Rylan Larsen; AAV-Ef1a- fDIO EYFP (#55641, in BLA) a gift from Karl Deisseroth and AAV5-Syn-FLEX-rc[ChrimsonR-tdTomato] (#62723, in BLA), a gift from Edward Boyden. AAV9-syn-FLEX-jGCaMP7f-WPRE (#104492) was a gift from Douglas Kim & GENIE Project.

### Stereotaxic surgery

At 6+ weeks of age, mice were anesthetized with 5% isoflurane. The hair over the incision cite was trimmed and the skin was prepped with alcohol and iodine. The animal was transferred to a stereotaxic frame (Kopf Instruments, Tujunga, CA) and kept under 1-2% isoflurane anesthesia. The skull surface was exposed via a midline sagittal incision and treated with the local anesthetic benzocaine (Medline Industries, Brentwood, TN). For each surgery, a 10uL microinjection syringe (Hamilton Co., Reno, NV) with a Micro4 pump controller (World Precision Instruments, Sarasota, FL) was guided by a motorized digital software (NeuroStar; Stoelting Co., Wood Dale, IL) to each injection coordinate. The following coordinates were used relative to Bregma, unless noted otherwise (in mm): PL (AP: 2.25, ML: ±0.40, DV: 2.09), DMT (AP: -1.00, ML: ±0.5, DV: 3.40), CeL (AP: -1.18, ML: ±2.77, DV: 4.80), BLA (AP: -1.35, ML: ±3.2-3.34, DV: 5.04-5.14), LEC (AP: 0.10, ML: ±4.28, DV: 4.42; relative to Lambda). For fiber photometry experiments, a fiber optic (Doric, 400 µm core, 0.66 NA) was implanted over the BLA injection site right after viral injection. All subjects received a 10 mg/kg ketoprofen (AlliVet, St. Hialeah, FL) injection as a perioperative analgesic, and additional post-operative treatment with ketoprofen was maintained for 48 h post-surgery.

### *Ex vivo* electrophysiology

Coronal brain sections were collected at 250μm using standard procedures. Mice were anesthetized using isoflurane, and transcardially perfused in an ice-cold/oxygenated (95% v/v O2, 5% v/v CO2) cutting solution consisting of (in mM): 93 N-Methyl-D-glucamine (NMDG), 2.5 KCl, 20 HEPES, 10 MgSO4. 7H2O, 1.2 NaH2PO4, 30 NaHCO3, 0.5 CaCl2. 2H2O, 25 glucose, 3 Na+-pyruvate, 5 Na+-ascorbate, and 5 N-acetylcysteine. The brain was subsequently dissected, hemisected, and sectioned using a vibrating LeicaVT1000S microtome (Leica Microsytems, Bannockburn, IL). The brain slices were then transferred to an oxygenated 34°C chamber filled with the same cutting solution for a 10 min recovery period. Slices were then transferred to a holding chamber containing a buffered solution consisting of (in mM): 92 NaCl, 2.5 KCl, 20 HEPES, 2 MgSO4 7H2O, 1.2 NaH2PO4, 30 NaHCO3, 2 CaCl2 2H2O, 25 glucose, 3 Na-pyruvate, 5 Na-ascorbate, 5 N-acetylcysteine and were allowed to recover for ≥30 min. For recording, slices were placed into a perfusion chamber where they were constantly exposed to oxygenated artificial cerebrospinal fluid (ACSF; 31-33°C) consisting of (in mM): 113 NaCl, 2.5 KCl, 1.2 MgSO4. 7H2O, 2.5 CaCl2. 2H2O, 1 NaH2PO4, 26 NaHCO3, 20 glucose, 3 Na+-pyruvate, 1 Na+-ascorbate, at a flow rate of 2.5-3ml/min.

Cells were visually identified from Ai14 reporter lines or virally injected animals under illumination from a series 120Q X-cite lamp at 40X magnification using an immersion objective in coordination with differential interference contrast microscopy (DIC). BLA neurons were either current clamped or voltage clamped in whole cell configuration using borosilicate glass pipettes (3-6MΩ) filled with intracellular solution, containing (in mM) either: 125 K+-gluconate, 4 NaCl, 10 HEPES, 4 Mg-ATP, 0.3 Na-GTP, and 10 Na-phosphocreatine, or 120 CsMeSO3, 2.8 NaCl, 5 TEA-Cl, 20 HEPES, 2.5 Mg-ATP and 0.25 Na-GTP (pH 7.30-7.35), respectively. In most cases pairs of neighboring pyramidal cell and interneuron were recorded simultaneously. Excitatory postsynaptic currents (EPSC) and inhibitory postsynaptic currents (IPSC) were recorded at -70mV and +10mV, respectively, using the CS-based internal solution. Experiments were generally conducted in a drug free ACSF. The following drugs have been applied when noted: Tetrodotoxin citrate (TTX, 1μM, Tocris Bioscience), 4-Aminopyridine (4AP, 100μM, Sigma-Aldrich), Gabazine (10μM, Sigma-Aldrich), Picrotoxin (50μM, Sigma-Aldrich), CNQX (20μM, Sigma-Aldrich), D(-)-2-Amino-5-Phosphonopentatonic acid (AP5, 50μM, Sigma-Aldrich), CGP-54626 (2μM, Sigma-Aldrich), Salvinorin B (SalB, 0.3μM, Tocris Bioscience).

All data was acquired using a Multiclamp 700B amplifier, Digidata 1440A A/D converter, Clampex version 10.6 software (Axon Instruments, Union City, CA), was sampled at 20kHz and low pass filtered at 1 kHz and analyzed in ClampFit 10 software (Molecular Devices, San Jose, CA). Optical stimulation was achieved by using a Mightex LED controller system with a 455nm blue LED and a 617nm red LED receiving and conveying TTL signals from Clampex protocols.

### Fear conditioning and extinction training

Behavioral fiber photometry experiments took place at least four weeks after stereotaxic surgeries allowing for GCaMP7f expression in the targeted BLA IN populations. Mice underwent extensive handling prior to training in order to eliminate basal fear expression. Two distinct contexts were used during the training protocol (Context A and B). Context A was used exclusively on fear conditioning day, and it consisted of vanilla scent, bright light, rectangular chamber with metal wiring floor for electric shock deliveries in a brightly lit room. Although Context B was presented in the same chamber as Context A, we used contrasting stimuli in order to help the differentiation between environments. Context B consisted of few drops of 2% acetic acid for olfactory cue, dark chamber with dim string lights, curved space created with a patterned plastic wall insert, white floor with clean bedding, within a dark room. Behavioral chamber was inside a Med Associates sound isolating box and Coulbourn Instruments was used for shock and sound deliveries using TTL signals initiated by FreezeFrame software.

Training started on day 1 in Context B referred to as habituation day (H). After 2-minute baseline mice were presented with eight 30s auditory tone (CS-: 400Hz, 75dB) unique to Context B with varying >30s inter-tone intervals (no tone). Mice displayed minimal freezing throughout the session (data not shown). The following day (day 2) fear conditioning took place in Context A. After 2-minute baseline 5 (initial SOMxAi14 cohort) to 6 distinct auditory tone-shock pairing was presented. Each tone (conditioned stimulus, CS+: 4000Hz, 75dB) lasted 30s and co-ended with a 1s electric foot shock (unconditioned stimulus, US: 0.45mA), with inter-tone intervals of 90s. For fear extinction training (day 3-5 or 6 if extinction has not developed faster) mice were placed in Context B and a combination of CS- (4-5, to test for tone-response specificity) and CS+ (8-14) tones were presented, each for 30s with varying >30s inter-tone intervals.

### Fiber photometry and analysis

Behavioral fiber photometry experiments took place at least four weeks after stereotaxic surgeries allowing for GCaMP7f expression in the targeted BLA IN populations. Mice were attached to a fiber optic patch cord (Doric, 400 µm core, 0.66 NA) and gently placed in the behavioral apparatus. Fiber photometry data was collected throughout the entire behavioral session using a rig with optical components from Doric lenses controlled by a real-time processor from Tucker Davis Technologies (TDT, RZ10x). TDT Synapse software was used for data acquisition. 465nm and 415nm (isosbestic control) LEDs were modulated at 210Hz and 330Hz, respectively. LED currents were adjusted to return a voltage between 75-85mV for each signal, were offset by 5mA, and were demodulated using a 10Hz lowpass frequency filter. For the analysis of the raw data, GuPPy ^43^, an open-source Python-based photometry data analysis pipeline, was used in order to visualize z-score traces and quantify peak z-score and area-under-the-curve (AUC) shown on figures using PSTH analysis. For CS+- and shock-related measurements 3s baseline prior to event was used, for all behavioral (start/stop freezing) transitions 1s baseline was used due to the high time-sensitivity of these events. Longer baselines shift the transients of single events resulting in less accurate data. Timestamps of start/stop freezing was extracted from FreezeFrame software (used for H-FC-E training) generated index files following careful manual determination of freezing thresholds for each mouse and inspection of the accuracy of chosen thresholds. Index files consisted of a string of 0/1 (motion/freezing) values assigned to time points. Timestamps of 01 (start freezing) and 10 (start moving) transitions were extracted. Since our behavioral (FreezeFrame) and photometry (Synapse) software was started manually at separate times, care needed to be taken to align the extracted index file timestamps to the photometry dataset. This time shift value could be assessed through the presence of known TTL signals (tone, shock) initiated by FreezeFrame to Synapse software. Additionally, a minimum duration criteria was applied for the 0/1 strings prior and after the transitions. We chose 1s-1s duration critera, which meant that only transitions where the animal was moving/freezing for at least 1s before it started freezing/moving for at least 1s were extracted. The reason for 1s is to eliminate potential artifacts yet keep most of the stable transitions in the analysis. Moreover, the timestamps were separated into CS-, CS+, and inter-tone interval (no tone) periods, and our analysis excluded the first 3-5 seconds of tone- and shock onset, -offset periods to eliminate confounding signals resulting from formerly uncovered phasic responses. A custom code was developed for timestamp extraction, which engages in a user-friendly interface (see Code availability, binary.py). These timestamps were visualized and analyzed by GuPPy. Another custom code was used for the calculation of overall ΔF/F values during motion and freezing separately, where data from CS and no tone periods were further differentiated after the elimination of the first 3-5 seconds of tone- and shock onset/offset. The code uses GuPPy generated ΔF/F and timestamp files together with the time-shifted FreezeFrame index file (see Code availability, dff.py). To confirm viral expression and fiber placement, mice were perfused, and brains were sectioned for histological verification using a fluorescent microscope (Keyence BZ-X800).

### Quantification and statistical analysis

Datasets were organized and quantified in Microsoft Excel and then transferred to GraphPad Prism 10 for generation of graphs and statistical analyses. For analysis of two groups, an unpaired or paired Student’s t test was used. For analysis of three or more groups across a single independent variable, an ordinary one-way ANOVA was used with Tukey post hoc multiple comparisons test between groups as noted in the figure legends. For analysis between two or more groups across two or more independent variables, a two-way ANOVA was used with a Sidak post hoc multiple comparisons test between groups as noted in the figure legends. Sample sizes were derived empirically and based on our previous experience with these assays. Significance was determined as p < 0.05 in all datasets.

## Code availability

Custom codes developed for this study are readily available on GitHub at https://github.com/sunilmut/NumPPy.

## ACKNOWLEDGMENTS

These studies were supported by NIH Grant MH11786 (S.P.) and a NARSAD Young Investigator Award (R.B.). We thank Megan Altemus, Keenan Johnson and Andrew Gaulden for excellent technical assistance throughout the completion of these studies.

## DECLERATIONS

S.M. is an employee of Microsoft Corporation (Redmond, WA), however volunteered his time for code development. All authors declare no financial conflicts of interest.

## AUTHOR CONTRIBUTIONS

R.B. conceived, designed, completed, and analyzed the experiments. S.M. contributed to code development as a volunteer, N.L. contributed to behavioral assays. R.B. and S.P. interpreted the results and prepared the manuscript.

---

**FIGURE S1.**
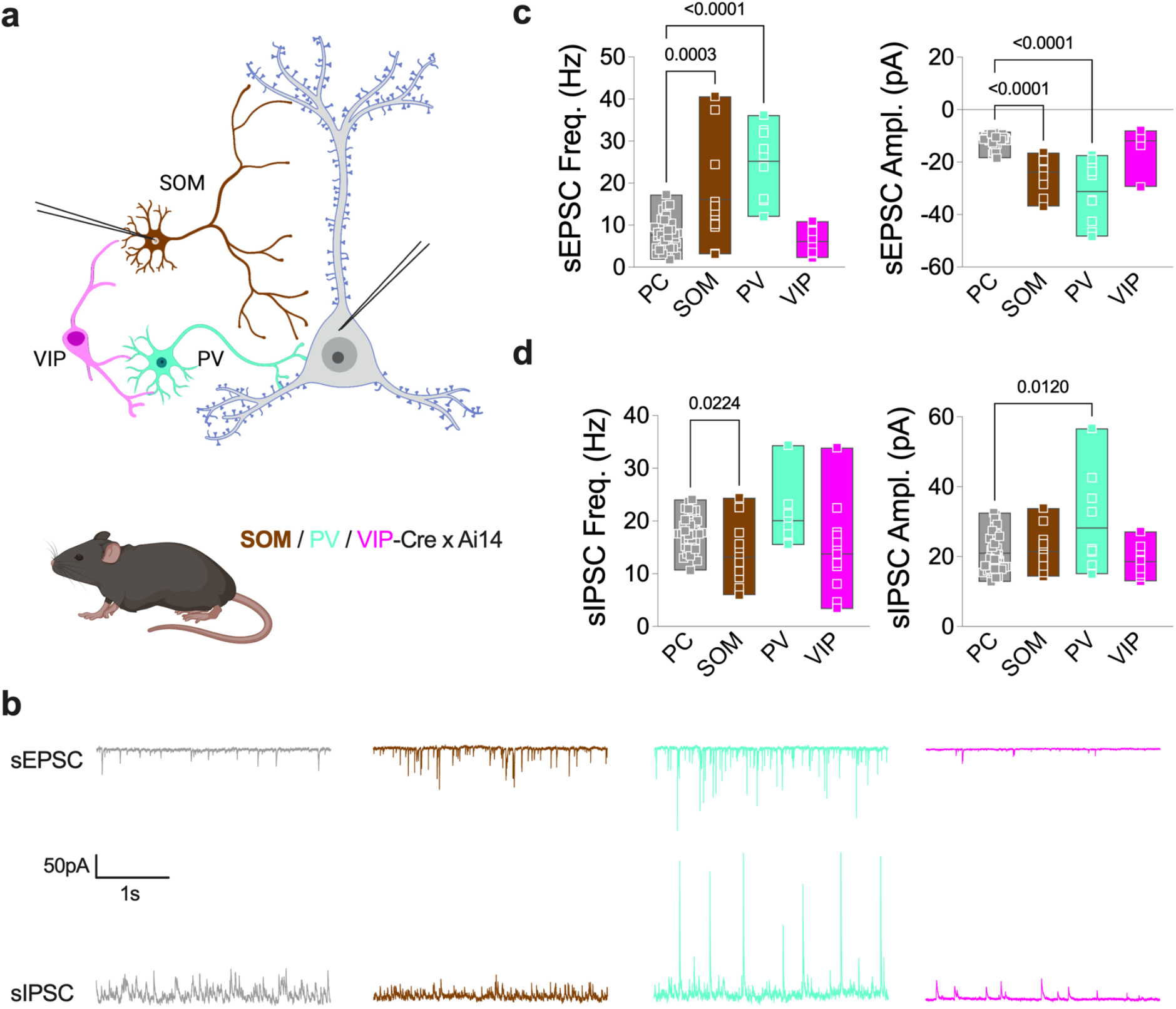
Analysis of spontaneous glutamatergic and GABAergic transmission onto BLA INs. a, Diagram of experimental approach. Whole cell voltage clamp recordings were obtained from SOM, PV or VIP INs and adjacent PCs. Glutamatergic and GABAergic transmissions were monitored in each cell at holding potentials of -70 mV and +10 mV, respectively. **b,** Representative traces of glutamatergic EPSCs and GABAergic IPSCs from PC (grey), SOM (brown), PV (green), and VIP (pink) IN. **c,** EPSC frequency and amplitude from BLA PCs, SOM, PV and VIP INs. **d,** IPSC frequency and amplitude from BLA PCs, SOM, PV, and VIP INs. Sample size (n cells, N mice); PCs (37,10), SOM (13, 3), PV (10,3), VIP (13, 4). **c-d,** P values and analysis via one-way ANOVA followed by Tukey post hoc tests (see **Table S1**).

**FIGURE S2.**
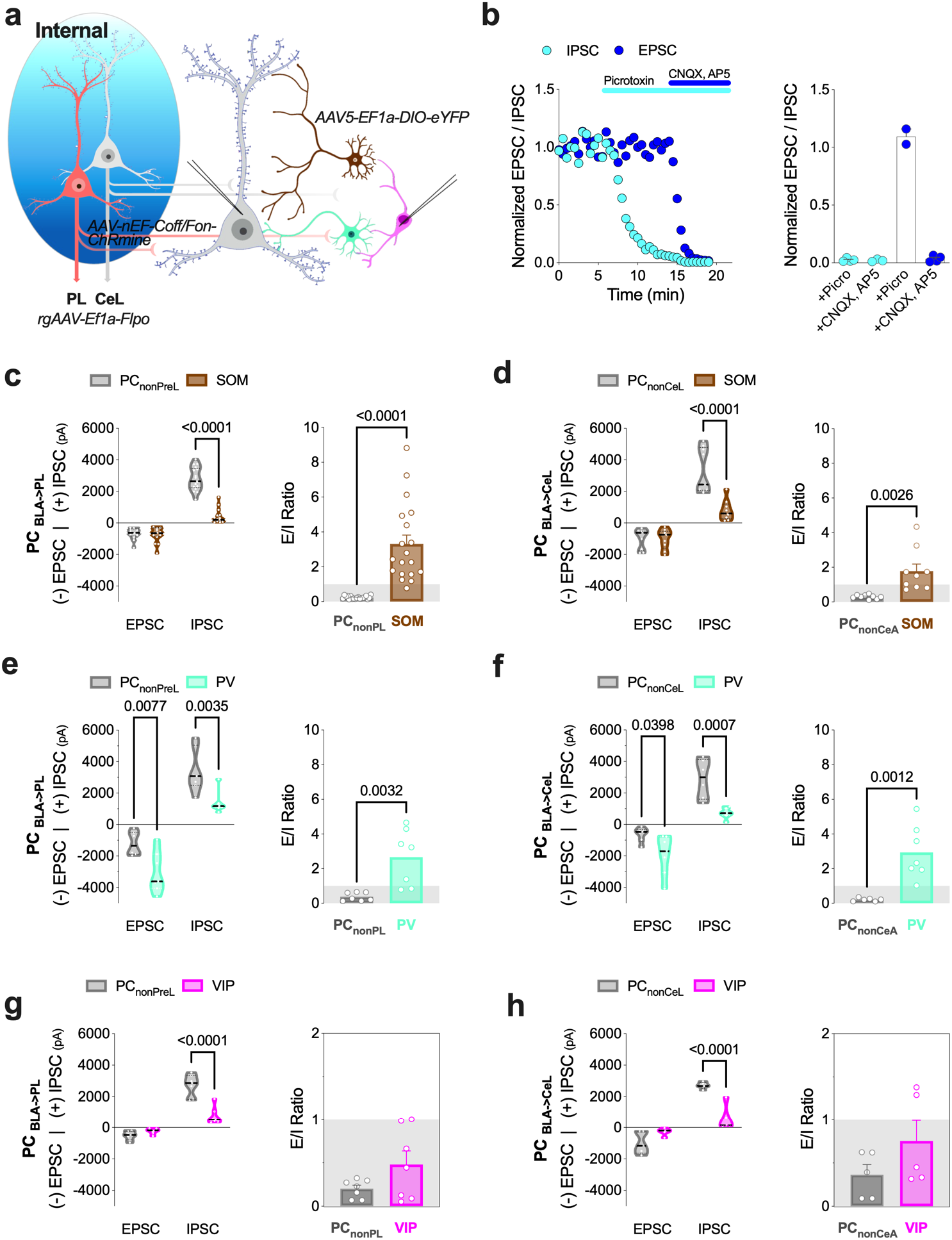
Effects of local BLA PC activation on glutamatergic and GABAergic transmission on different IN types. a, Diagram of experimental design. Local BLA PCs projecting to the CeL (PC _BLA -> CeL_) or PL (PC _BLA -> PL_) were transfected to express the excitatory opsin ChRmine and INs were labeled with Cre-dependent eYFP for visualization. Dual voltage- clamp recordings from identified INs and local non-labeled PCs were performed. **b,** Control experiments verifying isolated EPSCs and disynaptic IPSCs at -70mV and +10mV, respectively. **c-d,** PC _BLA-> PL_ and PC _BLA->CeL_ stimulation-elicited EPSC and IPSC amplitudes from SOM IN-PC pairs (left) and PC-IN excitation/inhibition (E/I) ratios (right). **e-f,** PC _BLA -> PL_ and PC _BLA -> CeL_ stimulation-evoked EPSC and IPSC amplitudes from PV IN-PC pairs and PC-IN E/I ratios. **g-h,** PC _BLA-> PL_ and PC _BLA-> CeL_stimulation-induced EPSC and IPSC amplitudes from VIP IN-PC pairs and PC-IN E/I ratios. Sample size (cell type, n cells, N mice); **c, e, g,** PC _BLA -> PL_ (SOM 17, 6; PC 17,6), (PV 7,3; PC 7,3), (VIP 7,4; PC 7,4); **d, f, h,** PC _BLA-> CeL_ (SOM 9,6; PC 9,6), (PV 7,2; PC 6,2), (VIP 6,3; PC 5,3). **c-h,** P values and data analysis via two-way ANOVA with Sidak post hoc test (left) and unpaired t-test (right) (see **Table S1**).

**FIGURE S3.**
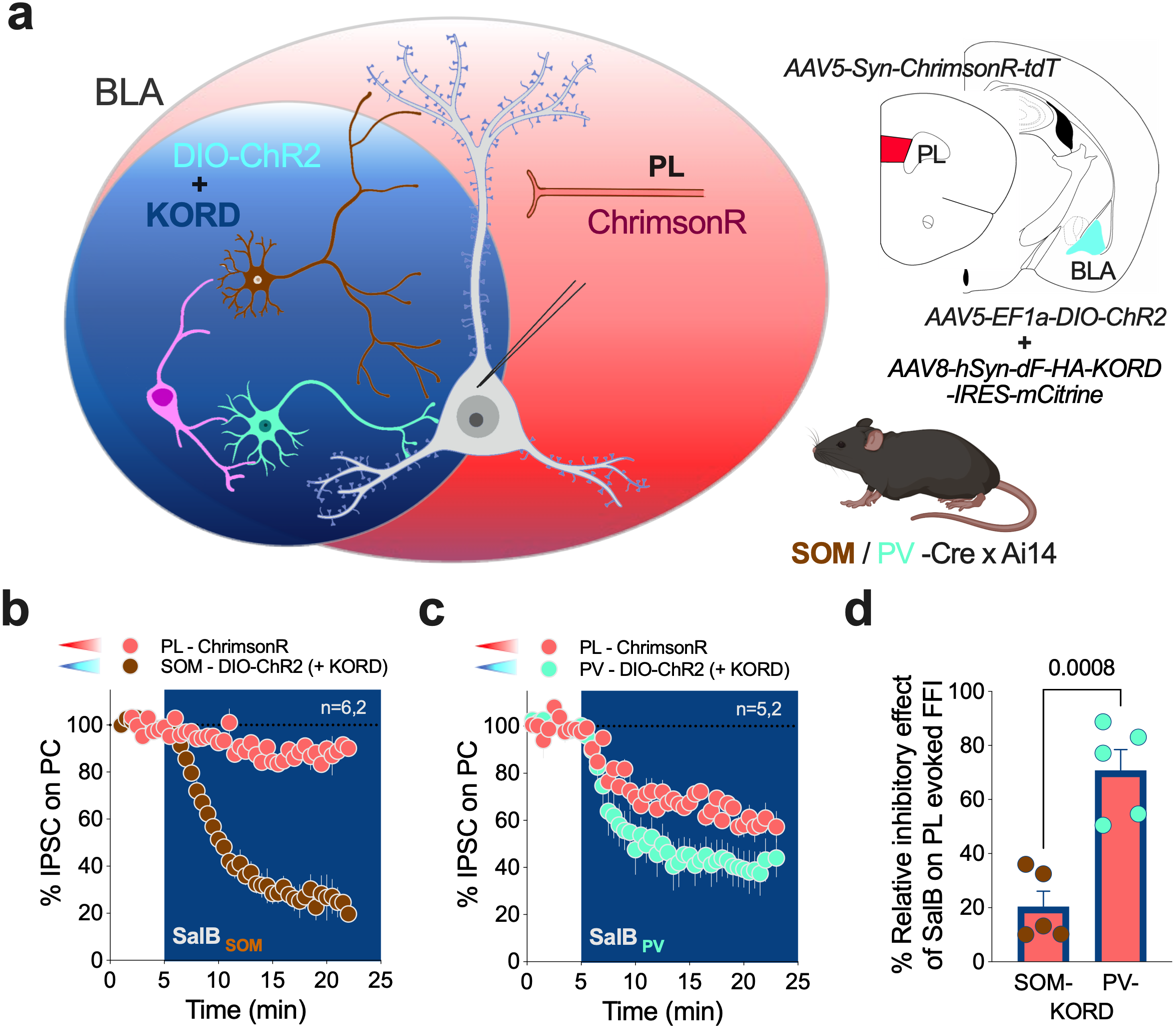
Analysis of feedforward inhibition onto SOM and PV INs upon PL afferent stimulation. a, Diagram of experimental design. ChrimsonR was expressed in PL neurons and Cre-dependent KORD and -ChR2 were expressed in distinct IN types. Patch clamp recordings were conducted from BLA PCs and feedforward inhibition (FFI) was monitored in response to red-light stimulation of PL inputs. Blue light activation of INs allowed for direct monitoring of IN-PC GABAergic transmission during application of Salvinorin B (SalB). **b,** SalB decreased SOM to PC GABAergic transmission but had minimal effect on feedforward inhibition elicited by PL stimulation. **c,** SalB decreased PV to PC GABAergic transmission and significantly reduced FFI calculated as a % of maximal SalB effect on direct IN to PC GABAergic transmission. Sample size (n cells, N mice); SOM (6,2) and PV (5,2). **d,** P value and analysis via unpaired t-test.

**FIGURE S4.**
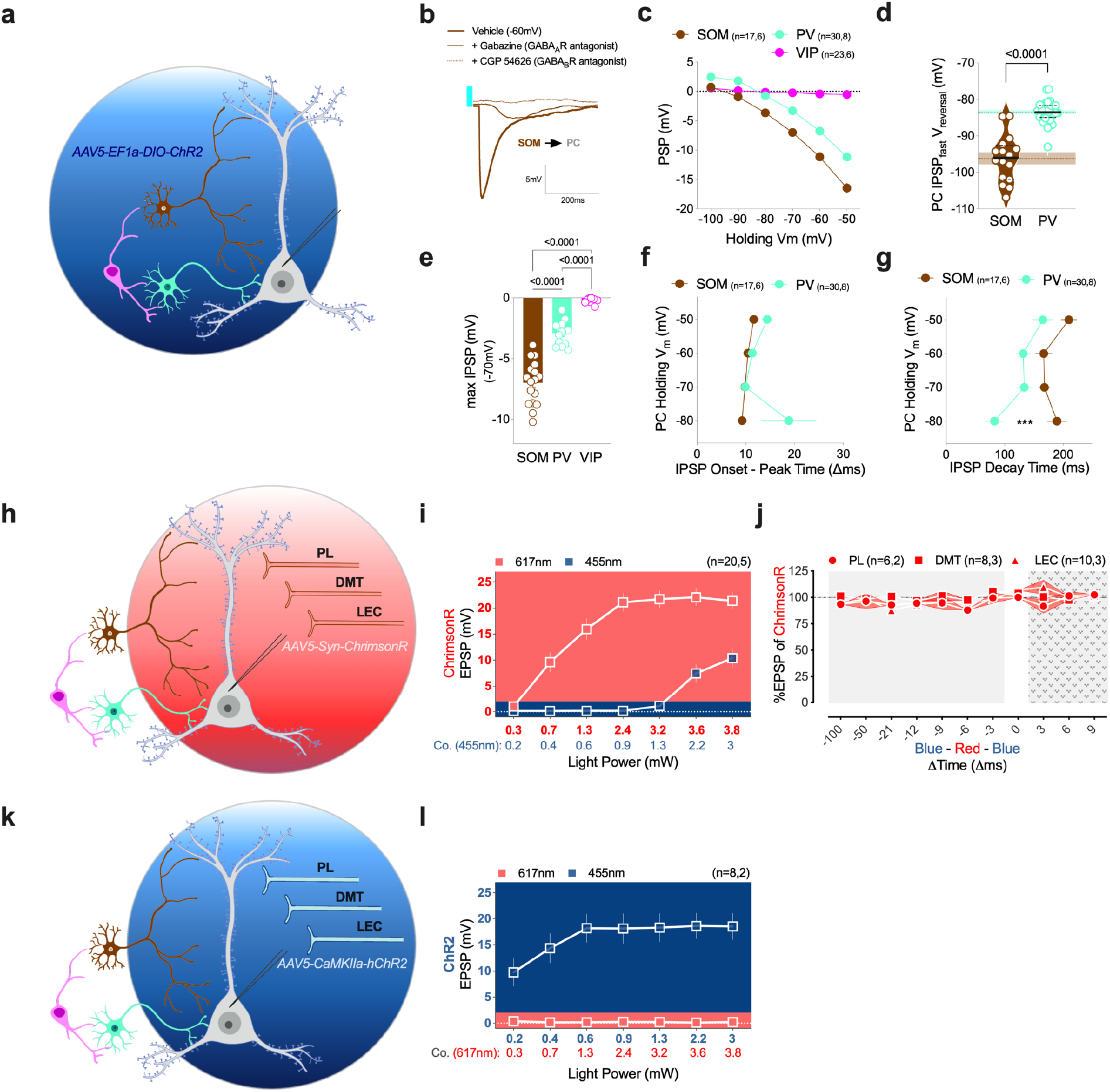
Properties of SOM and PV IN-mediated inhibition on PCs and control experiments supporting dual-opsin electrophysiology approaches. a, Diagram of experimental design. Cre-dependent ChR2 was expressed in either SOM or PV INs and whole cell patch clamp recordings were conducted from BLA PCs. **b,** Representative voltage trace from BLA PC following light activation of SOM INs showing prominent GABA_A_ receptor-mediated with some GABA_B_ receptor-mediated inhibitory components. **c,** BLA PC holding membrane potential and IPSP amplitude relationship evoked by SOM, PV and VIP IN stimulation. **d**, Reversal membrane potential of SOM and PV IN-mediated IPSPs. **e,** Maximal IPSP comparisons recorded from BLA PCs at a holding membrane potential of -70mV upon optical stimulation of SOM, PV or VIP INs. **f-g**, Onset-to-peak time and decay time of SOM and PV IN-mediated IPSPs at different holding membrane potentials. **h,** Control experiment where only ChrimsonR was expressed in PL, DMT or LEC without ChR2 in INs. **i,** Red light stimulation robustly evokes EPSPs in PCs while blue light does so only at light intensities above ∼1mW, thus in studies (Figure 3), blue light stimulation was kept below 0.5mV to avoid opsin cross-activation. **j,** EPSP amplitudes remained stable following red-light stimulation of PL, DMT or LEC afferents combined with blue-light (below 0.5mV) stimulation in varying time intervals in the absence of ChR2 expression. **k,** ChR2 was expressed in PL, DMT or LEC and patch clamp recordings were conducted from BLA PCs. **l,** Blue light stimulation of ChR2 terminals evoked robust EPSPs in BLA PCs whereas red light stimulation had no effect, even at high intensities. These data support specificity of dual-opsin approach used in main Figure 3. Sample size (n cells, N mice); **c-g** SOM (17,6), PV (30,8), VIP (23,6); **i** (20,5); **j,** PL (6,2), DMT (8,3), LEC (10,3); **l,** (8,2). **c**, P values and data analysis via 2way ANOVA followed by Tukey post hoc test; **d,** via unpaired t-test; **e,** analysis via one-way ANOVA followed by Tukey post hoc test; and **f-g,** via two-way ANOVA followed by Sidak post hoc test (see **Table S1**).

**FIGURE S5.**
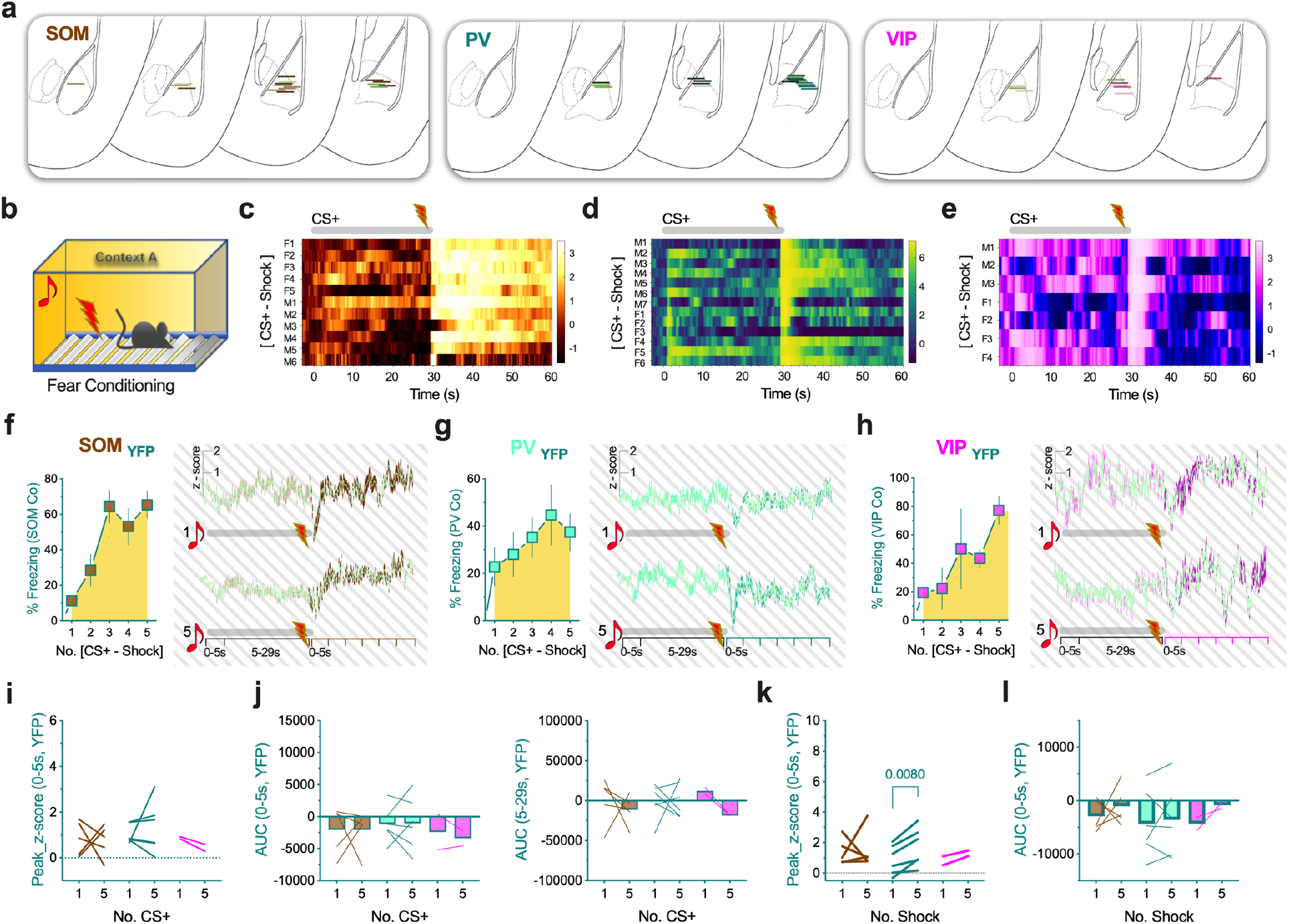
Control experiments related to IN fiber-photometry recordings during fear conditioning. a, BLA fiber implant placement for each mouse used in this study, bright green lines represent YFP-expressing control mice. Each mouse has undergone the complete habituation-fear conditioning-extinction behavioral paradigm. **b-e,** Heatmaps of z-score data averaged across all CS+-US pairings per mouse (F: female, M: male) during fear conditioning from SOM, PV, and VIP IN populations. **f-h,** Freezing data of control mice and control z-score traces of CS+-US pairing number 1 and 5 recorded from SOM, PV and VIP INs expressing YFP instead of GCaMP7f. **i,** Peak z-score following CS+ onset recorded from SOM-, PV- and VIP-YFP- expressing mice. **j,** AUC analysis (0-5s and 5-29s) in response to CS+ onset in SOM-, PV-, and VIP-YFP-expressing mice. **k,** Peak z-score following shock onset in SOM-, PV- and VIP-YFP- expressing mice. **l,** AUC analysis (0-5s) after shock onset in SOM-, PV-, and VIP-YFP-expressing mice. Sample size (n=number of mice; gender) **c-e,** SOM (n=11; 6M/5F), PV (n=13; 7M/6F), VIP (n=7; 3M/4F); **f-l,** for YFP controls (SOM n=5; 3M/2F), (PV n=6; 4M/2F), VIP (n=2M). **i-l,** P values and data analysis via paired t-test comparing CS+ or shock number 1 and 5 for each IN type.

**FIGURE S6.**
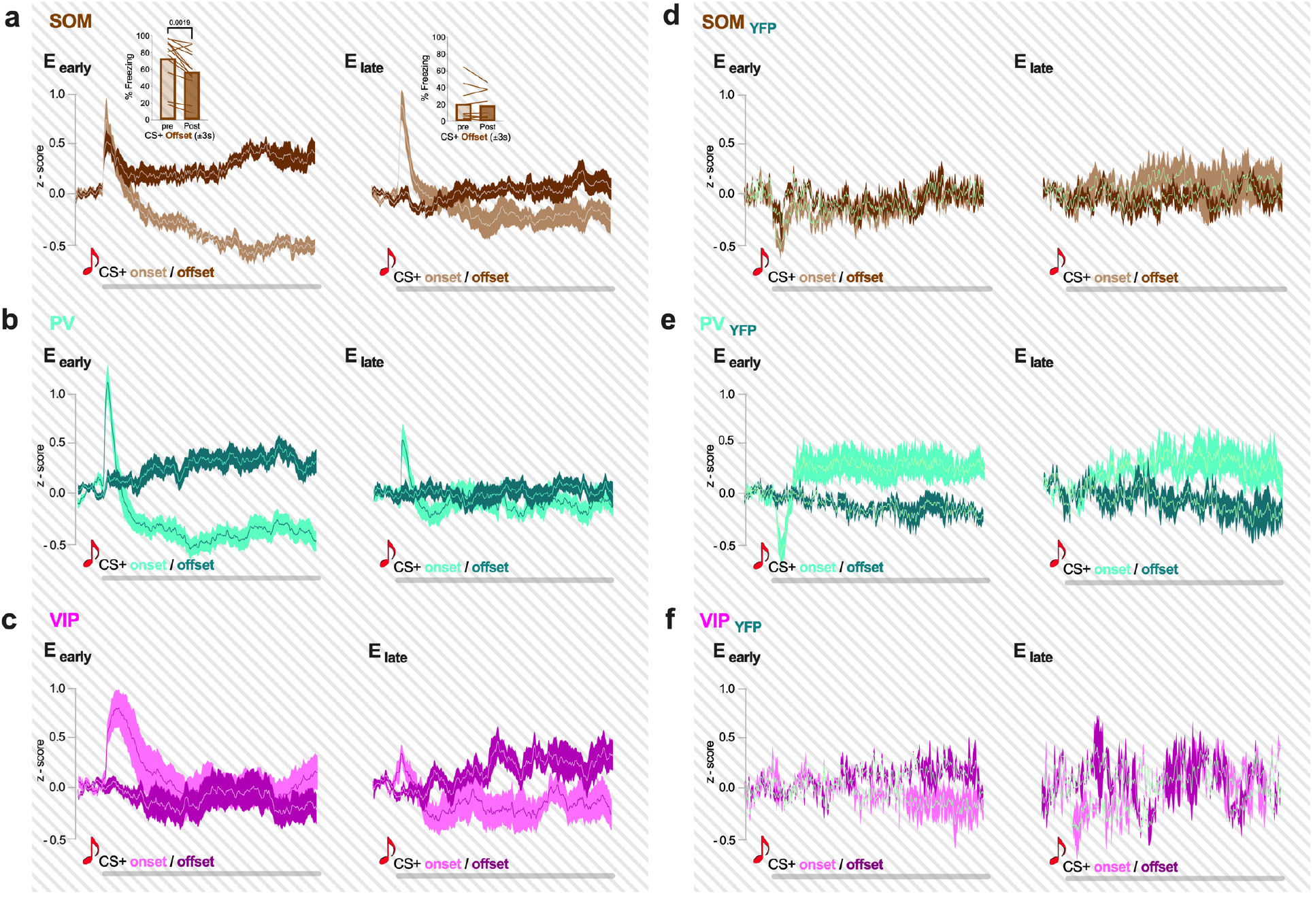
CS+ onset and offset responses during extinction in BLA IN types and related control data. a-c, Overlayed averaged z-score traces showing CS+ onset (light color) and offset (dark color) for SOM (**a**), PV (**b**), and VIP (**c**) INs during early and late phases of extinction (E_early_ and E_late_). Inset in **a,** shows freezing level changes between 3s prior and 3s post-CS+ offset during early and late extinction. **d-f,** Control data from SOM-, PV- and VIP-YFP expressing mice upon CS+ onset (light color) and offset (dark color) during early and late extinction. Z-score traces were first averaged per session, then across mice; traces represent mean ± SEM. Sample size: **a-c,** SOM (n=11; 6M/5F), PV (n=13; 7M/6F), VIP (n=7; 3M/4F); **d-f,** (SOM n=5; 3M/2F), (PV n=6; 4M, 2F), VIP (n=2M).

**FIGURE S7.**
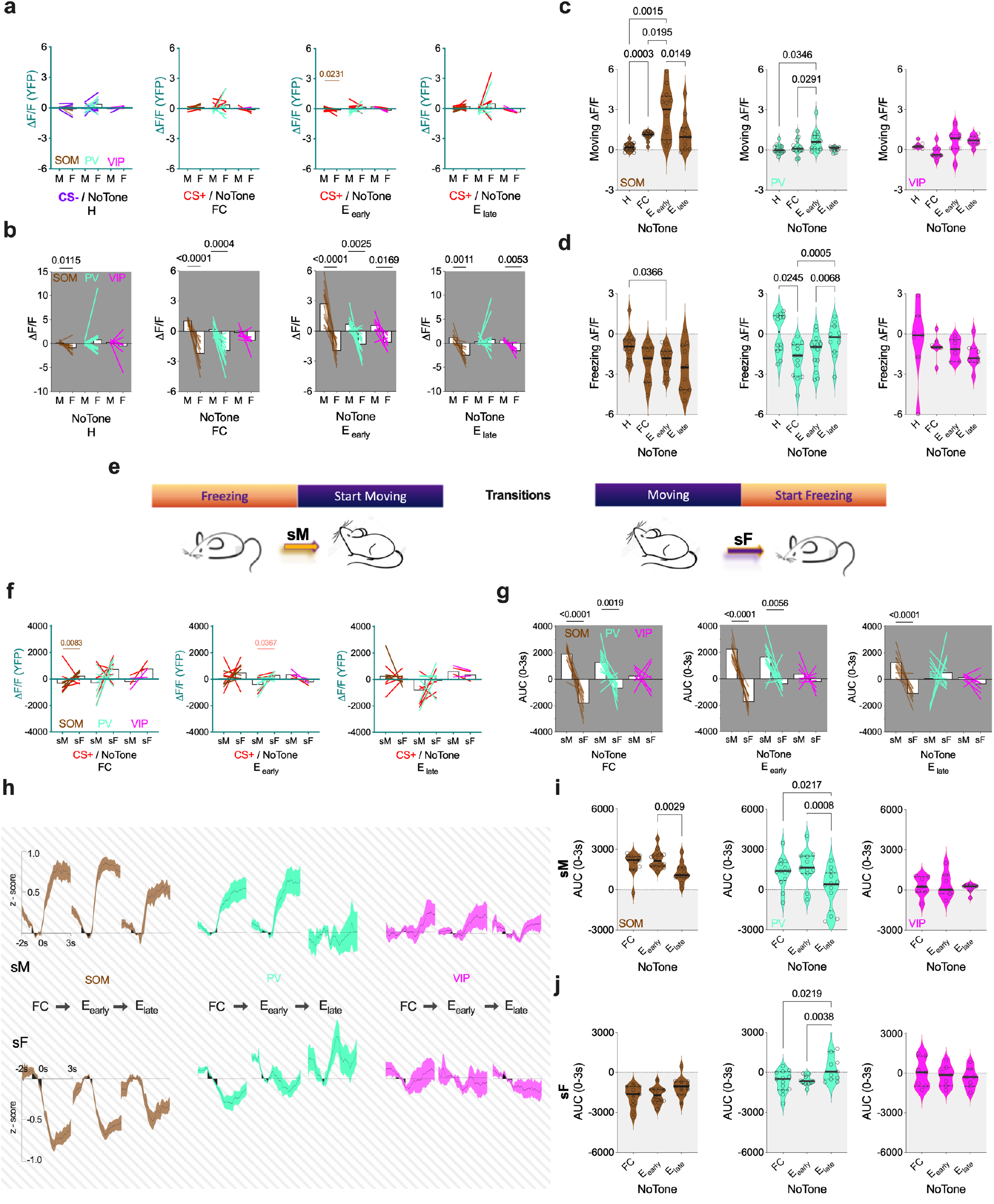
Behavioral state transition-related IN activity during inter-tone-intervals, and YFP control data. a, YFP-expressing control mice show no systematic changes in ΔF/F values as a function of movement or freezing either during CS presentations (purple and red lines) or during inter-tone-intervals (NoTone) across task days. **b,** Averaged ΔF/F values during bouts of motion (M) and freezing (F) on habituation (H), fear conditioning (FC), early- (E_early_) and late extinction (E_late_) days recorded from SOM (brown), PV (green) and VIP (pink) IN populations during inter-tone-intervals. **c,** Averaged ΔF/F values during bouts of motion compared across behavioral days for SOM, PV and VIP INs during inter-tone-intervals. **d,** Averaged ΔF/F values during bouts of freezing compared across behavioral days for SOM, PV and VIP INs during inter-tone-intervals. **e,** Diagram of behavioral state-transition analysis depicting dissection of transitions where mice start moving (sM) or start freezing (sF). Data analysis for habituation day was omitted due to low number of transitions per mouse. **f,** YFP-expressing control mice show no systematic changes in ΔF/F values between start moving and start freezing during CS+ presentations (red lines) or during inter-tone-intervals on separate task days. **g,** AUC (0-3s) values of start moving and start freezing transitions during inter-tone-intervals compared within IN types on each day for SOM, PV and VIP populations. **h,** Average z-score traces during behavioral state transitions for SOM, PV and VIP INs on fear conditioning, early and late extinction days. Z-score traces of transitions were first averaged per session, then across mice, traces represent mean ± SEM. Data were baselined to 1 second prior to behavioral state transition as shown in black. **i,** AUC (0-3s) values following motion onset (sM) during inter-tone-intervals compared across days for SOM, PV and VIP INs. **j,** AUC (0-3s) values following freezing onset (sF) during inter-tone-intervals compared across days for SOM, PV and VIP INs. Sample sizes; SOM (n=11; 6M/5F), PV (n=13; 7M/6F), VIP (n=7; 3M/4F). YFP-control sample sizes; (SOM n=5; 2M/3F), (PV n=6; 4M/2F), VIP (n=2M). **a, b, f, g,** P values and analysis via paired t-test comparing M to F and sM to sF for each IN type; **c, d, i, j,** analysis via one-way ANOVA followed by Tukey post hoc test (see **Table S1**).

**FIGURE S8.**
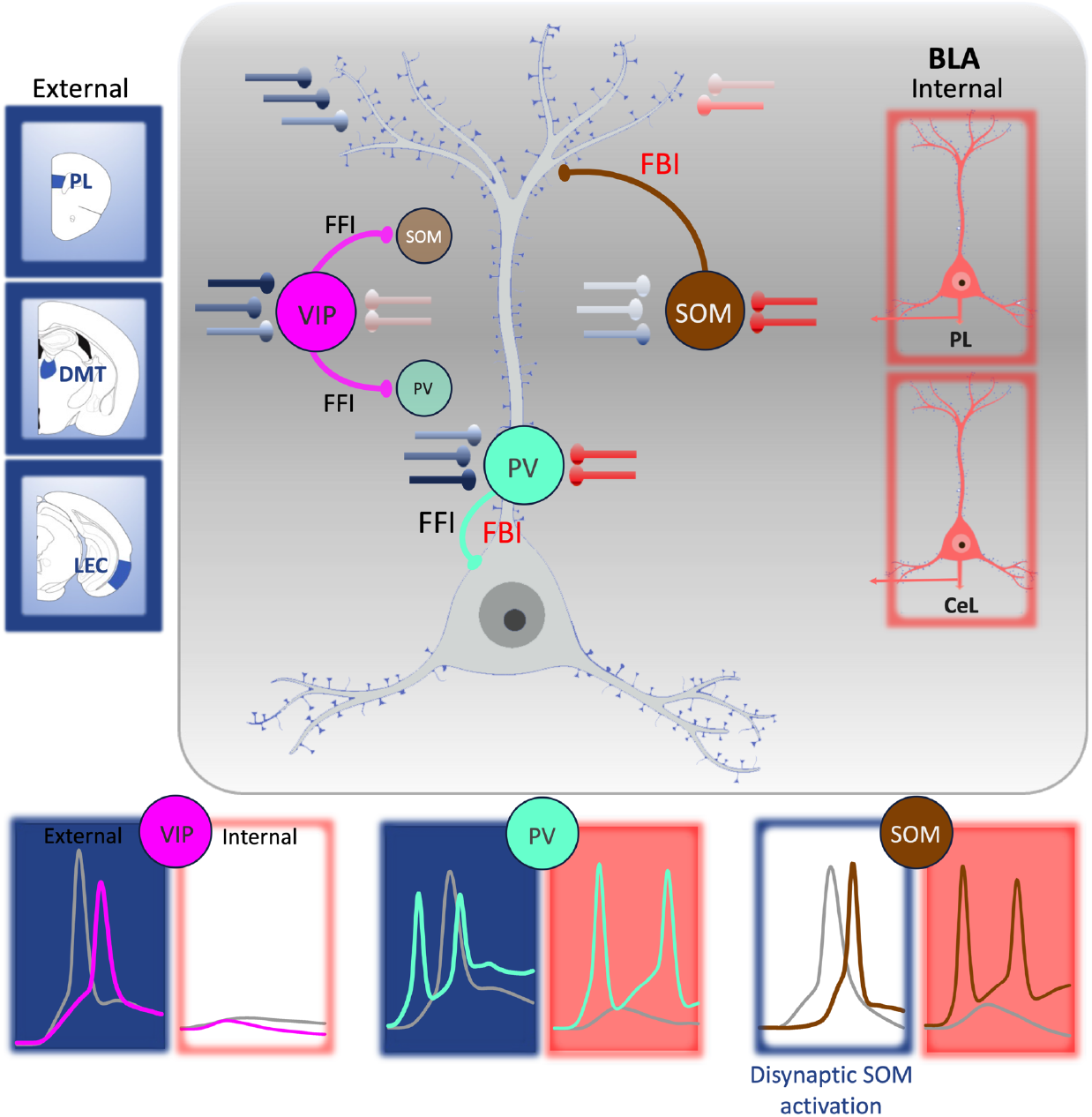
Overall model of BLA IN synaptic organization. VIP INs receive strong long- range afferents (blue) and generally do not inhibit BLA PCs, as a result provide feedforward inhibition (FFI) onto other INs. PV INs receive both long-range and local BLA inputs (red) and participate in both FFI and feedback inhibition (FBI), respectively. SOM INs get strong innervation from local PCs with minimal extrinsic afferents and thus participate in lateral or feedback inhibition of PCs. Weight of blue and red synapses reflect synaptic strength. For simplicity, only connections related to our study were depicted among BLA neurons.

**Table S1:**
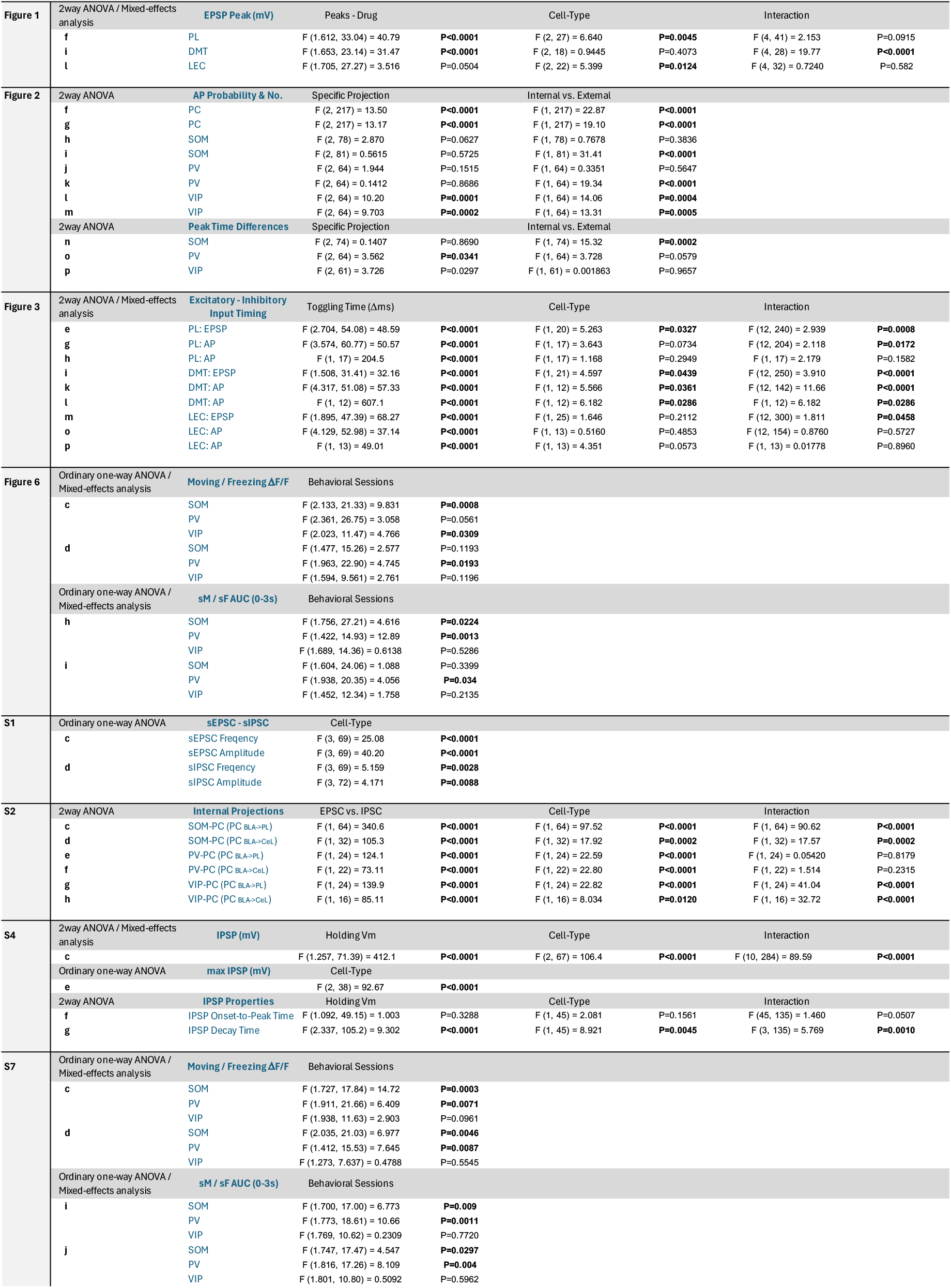

## Notes

### Competing Interest Statement

The authors have declared no competing interest.

## REFERENCES

1 Maren, S. Neurobiology of Pavlovian fear conditioning. Annu Rev Neurosci 24, 897–931 (2001). 10.1146/annurev.neuro.24.1.897

2 Fanselow, M. S. & Poulos, A. M. The neuroscience of mammalian associative learning. Annu Rev Psychol 56, 207–234 (2005). 10.1146/annurev.psych.56.091103.070213

3 Orsini, C. A. & Maren, S. Neural and cellular mechanisms of fear and extinction memory formation. Neurosci Biobehav Rev 36, 1773–1802 (2012). 10.1016/j.neubiorev.2011.12.014

4 Careaga, M. B. L., Girardi, C. E. N. & Suchecki, D. Understanding posttraumatic stress disorder through fear conditioning, extinction and reconsolidation. Neurosci Biobehav Rev 71, 48–57 (2016). 10.1016/j.neubiorev.2016.08.023

5 Lokshina, Y., Sheynin, J., Vogt, G. S. & Liberzon, I. Fear Extinction Learning in Posttraumatic Stress Disorder. Curr Top Behav Neurosci 64, 257–270 (2023). 10.1007/7854_2023_436

6 Bocchio, M., Nabavi, S. & Capogna, M. Synaptic Plasticity, Engrams, and Network Oscillations in Amygdala Circuits for Storage and Retrieval of Emotional Memories. Neuron 94, 731–743 (2017). 10.1016/j.neuron.2017.03.022

7 Johansen, J. P., Cain, C. K., Ostroff, L. E. & LeDoux, J. E. Molecular mechanisms of fear learning and memory. Cell 147, 509–524 (2011). 10.1016/j.cell.2011.10.009

8 Kim, J. J. & Jung, M. W. Neural circuits and mechanisms involved in Pavlovian fear conditioning: a critical review. Neurosci Biobehav Rev 30, 188–202 (2006). 10.1016/j.neubiorev.2005.06.005

9 McCullough, K. M. et al. Molecular characterization of Thy1 expressing fear-inhibiting neurons within the basolateral amygdala. Nat Commun 7, 13149 (2016). 10.1038/ncomms13149

10 Herry, C. et al. Switching on and off fear by distinct neuronal circuits. Nature 454, 600–606 (2008). 10.1038/nature07166

11 Senn, V. et al. Long-range connectivity defines behavioral specificity of amygdala neurons. Neuron 81, 428–437 (2014). 10.1016/j.neuron.2013.11.006

12 Kim, J., Pignatelli, M., Xu, S., Itohara, S. & Tonegawa, S. Antagonistic negative and positive neurons of the basolateral amygdala. Nat Neurosci 19, 1636–1646 (2016). 10.1038/nn.4414

13 Krabbe, S., Grundemann, J. & Luthi, A. Amygdala Inhibitory Circuits Regulate Associative Fear Conditioning. Biol Psychiatry 83, 800–809 (2018). 10.1016/j.biopsych.2017.10.006

14 Ehrlich, I. et al. Amygdala inhibitory circuits and the control of fear memory. Neuron 62, 757–771 (2009). 10.1016/j.neuron.2009.05.026

15 Lucas, E. K. & Clem, R. L. GABAergic interneurons: The orchestra or the conductor in fear learning and memory? Brain Res Bull 141, 13–19 (2018). 10.1016/j.brainresbull.2017.11.016

16 Cummings, K. A., Lacagnina, A. F. & Clem, R. L. GABAergic microcircuitry of fear memory encoding. Neurobiol Learn Mem 184, 107504 (2021). 10.1016/j.nlm.2021.107504

17 Wolff, S. B. et al. Amygdala interneuron subtypes control fear learning through disinhibition. Nature 509, 453–458 (2014). 10.1038/nature13258

18 Stujenske, J. M. et al. Prelimbic cortex drives discrimination of non-aversion via amygdala somatostatin interneurons. Neuron 110, 2258–2267 e2211 (2022). 10.1016/j.neuron.2022.03.020

19 Krabbe, S. et al. Adaptive disinhibitory gating by VIP interneurons permits associative learning. Nat Neurosci 22, 1834–1843 (2019). 10.1038/s41593-019-0508-y

20 Yau, J. O., Chaichim, C., Power, J. M. & McNally, G. P. The Roles of Basolateral Amygdala Parvalbumin Neurons in Fear Learning. J Neurosci 41, 9223–9234 (2021). 10.1523/JNEUROSCI.2461-20.2021

21 Majak, K. & Pitkanen, A. Activation of the amygdalo-entorhinal pathway in fear- conditioning in rat. Eur J Neurosci 18, 1652–1659 (2003). 10.1046/j.1460-9568.2003.02854.x

22 Giustino, T. F. & Maren, S. The Role of the Medial Prefrontal Cortex in the Conditioning and Extinction of Fear. Front Behav Neurosci 9, 298 (2015). 10.3389/fnbeh.2015.00298

23 Kirouac, G. J. Placing the paraventricular nucleus of the thalamus within the brain circuits that control behavior. Neurosci Biobehav Rev 56, 315–329 (2015). 10.1016/j.neubiorev.2015.08.005

24 Feldmeyer, D., Qi, G., Emmenegger, V. & Staiger, J. F. Inhibitory interneurons and their circuit motifs in the many layers of the barrel cortex. Neuroscience 368, 132–151 (2018). 10.1016/j.neuroscience.2017.05.027

25 Tremblay, R., Lee, S. & Rudy, B. GABAergic Interneurons in the Neocortex: From Cellular Properties to Circuits. Neuron 91, 260–292 (2016). 10.1016/j.neuron.2016.06.033

26 Smith, Y., Pare, J. F. & Pare, D. Differential innervation of parvalbumin-immunoreactive interneurons of the basolateral amygdaloid complex by cortical and intrinsic inputs. J Comp Neurol 416, 496–508 (2000).

27 Andrasi, T. et al. Differential excitatory control of 2 parallel basket cell networks in amygdala microcircuits. PLoS Biol 15, e2001421 (2017). 10.1371/journal.pbio.2001421

28 Guthman, E. M. et al. Cell-type-specific control of basolateral amygdala neuronal circuits via entorhinal cortex-driven feedforward inhibition. Elife 9 (2020). 10.7554/eLife.50601

29 Hajos, N. Interneuron Types and Their Circuits in the Basolateral Amygdala. Front Neural Circuits 15, 687257 (2021). 10.3389/fncir.2021.687257

30 Rhomberg, T. et al. Vasoactive Intestinal Polypeptide-Immunoreactive Interneurons within Circuits of the Mouse Basolateral Amygdala. J Neurosci 38, 6983–7003 (2018). 10.1523/JNEUROSCI.2063-17.2018

31 Lucas, E. K., Jegarl, A. M., Morishita, H. & Clem, R. L. Multimodal and Site-Specific Plasticity of Amygdala Parvalbumin Interneurons after Fear Learning. Neuron 91, 629–643 (2016). 10.1016/j.neuron.2016.06.032

32 Cummings, K. A. & Clem, R. L. Prefrontal somatostatin interneurons encode fear memory. Nat Neurosci 23, 61–74 (2020). 10.1038/s41593-019-0552-7

33 Mineur, Y. S. et al. ACh signaling modulates activity of the GABAergic signaling network in the basolateral amygdala and behavior in stress-relevant paradigms. Mol Psychiatry 27, 4918–4927 (2022). 10.1038/s41380-022-01749-7

34 Kim, T. et al. Activated somatostatin interneurons orchestrate memory microcircuits. Neuron 112, 201–208 e204 (2024). 10.1016/j.neuron.2023.10.013

35 Piantadosi, S. C. et al. Holographic stimulation of opposing amygdala ensembles bidirectionally modulates valence-specific behavior via mutual inhibition. Neuron 112, 593–610 e595 (2024). 10.1016/j.neuron.2023.11.007

36 Kyriazi, P., Headley, D. B. & Pare, D. Different Multidimensional Representations across the Amygdalo-Prefrontal Network during an Approach-Avoidance Task. Neuron 107, 717–730 e715 (2020). 10.1016/j.neuron.2020.05.039

37 Kyriazi, P., Headley, D. B. & Pare, D. Multi-dimensional Coding by Basolateral Amygdala Neurons. Neuron 99, 1315–1328 e1315 (2018). 10.1016/j.neuron.2018.07.036

38 Fustinana, M. S., Eichlisberger, T., Bouwmeester, T., Bitterman, Y. & Luthi, A. State- dependent encoding of exploratory behaviour in the amygdala. Nature 592, 267–271 (2021). 10.1038/s41586-021-03301-z

39 Grundemann, J. et al. Amygdala ensembles encode behavioral states. Science 364 (2019). 10.1126/science.aav8736

40 Pantazopoulos, H., Lange, N., Hassinger, L. & Berretta, S. Subpopulations of neurons expressing parvalbumin in the human amygdala. J Comp Neurol 496, 706–722 (2006). 10.1002/cne.20961

41 Gouwens, N. W. et al. Integrated Morphoelectric and Transcriptomic Classification of Cortical GABAergic Cells. Cell 183, 935–953 e919 (2020). 10.1016/j.cell.2020.09.057

42 Nakashima, M., Ikegaya, Y. & Morikawa, S. Genetic labeling of axo-axonic cells in the basolateral amygdala. Neurosci Res 178, 33–40 (2022). 10.1016/j.neures.2022.02.002

43 Sherathiya, V. N., Schaid, M. D., Seiler, J. L., Lopez, G. C. & Lerner, T. N. GuPPy, a Python toolbox for the analysis of fiber photometry data. Sci Rep 11, 24212 (2021). 10.1038/s41598-021-03626-9

